# Plastoquinone redox status influences carboxysome integrity via a RpaA- and ROS-dependent regulatory network

**DOI:** 10.1101/2025.01.24.634715

**Authors:** María Santos-Merino, Lauri Nikkanen, Emmanuel J. Kokarakis, Yagut Allahverdiyeva, Daniel C. Ducat

**Affiliations:** MSU-DOE Plant Research Laboratory, Michigan State University, East Lansing, MI, United States, 48824; Laboratory of Molecular Plant Biology, Department of Life Technologies, University of Turku, Turku, Finland, 20014; Department of Microbiology and Molecular Genetics, Michigan State University, East Lansing, MI, United States, 48824; Department of Biochemistry and Molecular Biology, Michigan State University, East Lansing, MI, United States, 48824

## Abstract

Carboxysomes are bacterial microcompartments that encapsulate Rubisco and are a core component of the cyanobacterial carbon concentration mechanism (CCM). While carboxysome number, size and spatial organization are observed to vary in different environmental conditions (CO_2_, light, temperature, light quality), molecular mechanisms underlying this potentially adaptive process remain elusive. Herein, we observed that mutants of the circadian rhythm/metabolism factor, Regulator of Phycobilisome Associated A (RpaA), exhibit a striking breakdown of carboxysomes under certain environmental conditions. We find that growth conditions leading to overreduction of the plastoquinone (PQ) pool (mixotrophic growth, high irradiance, or chemical inhibition of electron transfer from PQ to the cytochrome *b_6_f* complex) are accompanied by elevated generation of reactive oxygen species (ROS), and correlate with carboxysome breakdown. Carboxysome breakdown is reversed by environmental conditions or chemical inhibitors that prevent PQ overreduction and accompanying ROS generation. Taken together, our data supports a novel link between the redox status of the PQ pool and carboxysome status and/or integrity. Our results have implications for fundamental understanding of cyanobacterial energy balancing pathways and may indicate new research directions for understanding how the carboxysome is remodeled in response to changing environments.

## 1. Introduction

Cyanobacteria and other photosynthetic organisms employ a variety of adaptive pathways to contend with variability in light, which fluctuates in predictable (*e.g.*, diel cycles, seasonal variations) and stochastic (*e.g.*, environmental shading, light flecks) patterns (Labiosa et al., 2006). High irradiance or spikes in illumination can result in an over-reduced photosynthetic electron transport chain and lead to the generation of toxic reactive oxygen species (ROS) (Foyer, 2018; Calzadilla and Kirilovsky, 2020). Photosynthetic organisms therefore possess a variety of regulatory mechanisms to coordinate the upstream processes of photochemical activity with downstream metabolic processes energetic demands to achieve a balance between light harvesting and energy utilization. In photosynthetic linear electron flow, electrons extracted from water are transferred from photosystem II (PSII) to photosystem I (PSI) via a series of electron carriers, including the membrane soluble plastoquinone (PQ) pool. While the ratio of proportional increase in ATP/NADPH requirements can create an energy imbalance, leading to the reduction or oxidation of the intersystem electron transport chain, and more specifically, the plastoquinone (PQ) pool (Fujita et al., 1987; Sunil et al., 2013; Fan et al., 2021). State transitions (Calzadilla and Kirilovsky, 2020) and photoprotective systems (such as orange carotenoid protein or flavodiiron proteins) (Kirilovsky and Kerfeld, 2016; Nikkanen et al., 2021) are well-studied processes by which cyanobacteria attempt to rebalance the redox status of the pETC in the short term. While the formation of ROS is typically discussed as a dangerous byproduct of unbalanced photosynthetic activity, there are also well-established signaling roles for some of these molecules, especially hydrogen peroxide (H_2_O_2_) (Mironov et al., 2019).

In addition to short-term changes in light availability, day/night cycles introduce irradiance changes that are anticipated by the well-characterized cyanobacterial circadian rhythm machinery (Dong et al., 2010; Yang et al., 2010; Pattanayak et al., 2014; Cohen and Golden, 2015; Diamond et al., 2015; Martins et al., 2018; Taton et al., 2020). The cyanobacterial circadian clock consists of a core oscillator, KaiC, which undergoes cyclic rounds of phosphorylation and dephosphorylation that are programmed by upstream regulators connected to light reactions via the PQ pool and has output functions controlled by the two-component system proteins SasA and RpaA (Nakajima et al., 2005; Ivleva et al., 2006; Takai et al., 2006; Kim et al., 2012; Kim et al., 2020). RpaA is a transcription factor that acts as a master regulator of the circadian clock outputs, by driving global rhythms of gene expression and gating of cell division (Dong et al., 2010; Markson et al., 2013). RpaA was first discovered in *Synechocystis* sp. PCC 6803, as an OmpR-type response regulator that regulates the energy transfer from phycobilisomes to PSI and PSII (Ashby and Mullineaux, 1999). Recently, other functions have been subscribed to RpaA beyond its roles in entraining the circadian clock and controlling clock output (Diamond et al., 2015; Iijima et al., 2015; Diamond et al., 2017; Puszynska and O’Shea, 2017). For example, it has been proposed that RpaA may be more directly involved in regulating enzymes of core metabolism to influence carbon partitioning (*i.e.*, towards glycogen storage vs. downstream metabolism) (Puszynska and O’Shea, 2017).

The carboxysome is an essential complex of the cyanobacterial carbon concentration mechanism (CCM) and is also dynamically regulated in response to changing light and other environmental conditions (Burnap et al., 2013). Carboxysomes are bacterial microcompartments comprised of a proteinaceous shell that encapsulates Rubisco. The primary function of the carboxysome is to maximize the carbon-fixing capacity of Rubisco by defining a microenvironment that maximizes inorganic carbon availability and simultaneously minimizes photorespiratory flux (Long et al., 2007). Carboxysomes respond to environmental changes by adjusting in number, size, and spatial organization based on CO_2_, light availability, redox state, temperature, and light quality (Sun et al., 2016; Rohnke et al., 2018; Sun et al., 2019; Rillema et al., 2021; Lucius and Hagemann, 2024). Their positioning and carbon fixation capacity also vary with diurnal cycles and carbon demand (Sun et al., 2020; Singh et al., 2022). For example, in the model cyanobacterium *Synechococcous elongatus* PCC 7942 (*S. elongatus*), an increase in irradiance is generally correlated with increased carboxysome number (Sun et al., 2016), suggestive of a regulatory link between the availability of light energy and cellular investment in carbon fixation machinery. Similarly, we and others have recently shown that *S. elongatus* engineered to have a higher metabolic “demand” via the expression of heterologous metabolism also exhibit an increase in carboxysome number (Santos-Merino et al., 2021a; Singh et al., 2022). Therefore, the “upstream” processes of photosynthetic light reactions appear to be integrated with the “downstream” energy demands of the cell for carbon fixation and metabolism, although regulatory mechanisms that accomplish this are poorly understood (Santos-Merino et al., 2021a).

We recently conducted a screen of all known two-component regulatory proteins *S. elongatus* with the goal of identifying potential cyanobacterial networks important for achieving energy balance between photosynthetic energy harvesting and integrated metabolic demand (Santos-Merino et al., 2024). Briefly, our approach used a heterologous metabolic pathway (sucrose secretion) that can be experimentally activated to significantly draw upon primary products of photosynthesis. Activation of the sucrose secretion pathway has been previously shown to lead to a range of photosynthetic changes, including increased carboxysome number, increased CO_2_ fixation rates, increased oxygen evolution, and reduced acceptor side limitation of PSI activity (Abramson et al., 2016; Santos-Merino et al., 2021b; Singh et al., 2022). Our screen for regulatory proteins involved in this process implicated RpaA, ManS, CikB, and NblS as leading factors important in coupling changes in metabolic demand in the cell to upstream enhancements in the flux photosynthetic processes. Herein, we report potential mechanisms that link RpaA function to the control of the organization of the carboxysome in response to ROS signals derived from an over-reduced pETC. Our results have implications for fundamental understanding of cyanobacterial energy balancing pathways and may provide insight into how the carboxysome is remodeled in response to changing environments and the utilization of central carbon metabolic intermediates.

## 2. Materials and methods

### 2.1. Strains and culture conditions

*S. elongatus* cultures were grown in BG11 medium supplemented with 1 g L^−1^ HEPES to a final pH of 8.3 with NaOH. Flasks were cultured in a Multitron incubator (Infors HT) at 32 °C under ambient air CO_2_ or supplemented with 2% CO_2_ with ∼150 μmol photons m^−2^ s^−1^ of light provided by Sylvania 15 W Gro-Lux fluorescent bulbs and shaken at 150 rpm. Cultures were back-diluted daily to an OD_750_ of 0.3 and acclimated to the medium/irradiance for at least 3 days prior to experiments or isopropyl-β-D-thiogalactoside (IPTG) induction. Where appropriate, 1 mM IPTG was added to induce *cscB* and *sps* gene expression. Erythromycin (Em; 100 μg mL^−1^), chloramphenicol (Cm; 25 μg mL^−1^), and spectinomycin (Sp; 100 μg mL^−1^) were used to maintain *cscB-sps*-, *cscB*-, and *rbcS-mNG*-containing cells, respectively. Kanamycin (Kn; 12.5 μg mL^−1^) was used to maintain the *rpaA* inactivation mutant Δ*rpaA*. In all cases, antibiotic selection was removed prior to conducting any of the reported experiments to minimize any unintended effects. When indicated, cultures were grown in the presence of photosynthesis inhibitors, including DCMU (20 µM) or DBMIB (10 µM). All strains used in this study are listed in Table 1.

**Table 1.** Cyanobacterial strains used in this study.

| Strain | Relevant genotype and phenotype<sup>a</sup> | Plasmids used to generate this strain | Source or reference |
| --- | --- | --- | --- |
| CscB-SPS<sup>export</sup> | *S. elongatus* PCC 7942 with P<sub>trc</sub>::*cscB* and P<sub>trc</sub>::*sps* integrated at NS3; Em<sup>r</sup> |  | (Santos-Merino et al., 2024) |
| CscB-SPS<sup>export</sup>/Δ*rpaA* | *S. elongatus* PCC 7942 with P<sub>trc</sub>::*cscB* and P<sub>trc</sub>::*sps* integrated at NS3, and with Δ*rpaA*::*aphI*; Em<sup>r</sup> Kn<sup>r</sup> |  | (Santos-Merino et al., 2024) |
| CscB-SPS<sup>export</sup>/RbcS-mNG | *S. elongatus* PCC 7942 with P<sub>trc</sub>::*cscB* and P<sub>trc</sub>::*sps* integrated at NS3, and with P<sub>rbcS</sub>::*rbcS-mNG* integrated at NS1; Em<sup>r</sup> Sp<sup>r</sup> | pRbcS-mNG | This work |
| CscB-SPS<sup>export</sup>/RbcS-mNG/Δ*rpaA* | *S. elongatus* PCC 7942 with P<sub>trc</sub>::*cscB* and P<sub>trc</sub>::*sps* integrated at NS3, with P<sub>rbcS</sub>::*rbcS-mNG* integrated at NS1, and with Δ*rpaA*::*aphI*; Em<sup>r</sup> Sp<sup>r</sup> Kn<sup>r</sup> | pRbcS-mNG<br>pMD19-T-0095 | This work |
| CscB<sup>import</sup> | *S. elongatus* PCC 7942 with P<sub>trc</sub>::*cscB* integrated at NS3; Cm<sup>r</sup> |  | (Ducat et al., 2012) |
| CscB<sup>import</sup>/RbcS-mNG | *S. elongatus* PCC 7942 with P<sub>trc</sub>::*cscB* integrated at NS3, and with P<sub>rbcS</sub>::*rbcS-mNG* integrated at NS1; Cm<sup>r</sup> Sp<sup>r</sup> | pRbcS-mNG | This work |
| CscB<sup>import</sup>/RbcS-mNG/Δ*rpaA* | *S. elongatus* PCC 7942 with P<sub>trc</sub>::*cscB* integrated at NS3, with P<sub>rbcS</sub>::*rbcS-mNG* integrated at NS1, and with Δ*rpaA*::*aphI*; Cm<sup>r</sup> Sp<sup>r</sup> Kn<sup>r</sup> | pRbcS-mNG<br>pMD19-T-0095 | This work |
| CscB<sup>import</sup>/RbcS-mNG | *S. elongatus* PCC 7942 with P<sub>trc</sub>::*cscB* integrated at NS3, and with the native copy of *mcdB* replaced with *mNG-rbcS*; Cm<sup>r</sup> Kn<sup>r</sup> | pmNG-McdB | This work |
| CscB<sup>import</sup>/mNG-McdB/Δ*rpaA* | *S. elongatus* PCC 7942 with P<sub>trc</sub>::*cscB* integrated at NS3, with the native copy of *mcdB* replaced with *mNG-rbcS* and with Δ*rpaA*::*aacCI*; Cm<sup>r</sup> Kn<sup>r</sup> Gm<sup>r</sup> | pmNG-McdB<br>pMD19-T-0095 | This work |

<sup>a</sup>Em<sup>r</sup>, erythromycin resistance; Kn<sup>r</sup>, kanamycin resistance; Sp<sup>r</sup>, spectinomycin resistance; Cm<sup>r</sup>, chloramphenicol resistance; NS1, neutral site 1; NS3, neutral site 3.

### 2.2. Strain construction

*S. elongatus* with genomically integrated copies of *cscB* and *sps* under an IPTG-inducible promoter was previously obtained (Santos-Merino et al., 2024), and *S. elongatus* with genomically integrated copies of *cscB* under an IPTG-inducible promoter was previously described (Ducat et al., 2012). Genomic loci encoding *rpaA* gene was disrupted by inserting a kanamycin resistance cassette (Qiao et al., 2019). To allow visualization of changes in carboxysome organization, we integrated a fluorescent reporter fused to the small subunit of Rubisco (RbcS-mNG) in NS1 by modifying a plasmid previously published (Sakkos et al., 2021). Plasmid details are reported in Table 2.

**Table 2.** Plasmids used in this study.

| Plasmid | Relevant genotype and phenotype<sup>a</sup> | Source or reference |
| --- | --- | --- |
| pAM2314-RbcS-mNG | pAM2314 with $P_{rbcLS}::rbcS$ fused to mNG for neutral site 1 integration; Cm<sup>r</sup> | (Sakkos et al., 2021) |
| pRbcS-mNG | Derivative of pAM2314-RbcS-mNG modified to replace *cat* gene conferring chloramphenicol resistance by *aadA* gene conferring spectinomycin resistance; Sp<sup>r</sup> | This work |
| pMD19-T-0095 | Plasmid to replace the *rpaA* gene with kanamycin resistance cassette (*aphI*); Ap<sup>r</sup> Kn<sup>r</sup> | (Qiao et al., 2019) |
| pmNG-McdB | Plasmid to replace the endogenous copy of *mcdB* by a version fused to mNG; Kn<sup>r</sup> | (MacCready et al., 2018) |

<sup>a</sup>Cm<sup>r</sup>, chloramphenicol resistance; Sp<sup>r</sup>, spectinomycin resistance; Ap<sup>r</sup>, ampicillin resistance; Kn<sup>r</sup>, kanamycin resistance.

### 2.3. Sucrose quantification

Secreted sucrose was quantified from supernatants using the Sucrose/D-Glucose Assay Kit (K-SUCGL; Megazyme).

### 2.4. Pigment determination

Chl*a* was extracted from cell pellets by incubation in 100% methanol for 30 min at 4 °C. Chl*a* concentration was estimated by the spectrophotometric method described previously (Porra et al., 1989).

### 2.5. Fluorescence measurements

Apparent quantum yield of PSII (Φ_II_) measurements were performed on a custom-built fluorimeter/spectrophotometer as described previously (Santos-Merino et al., 2021b). Briefly, samples containing cyanobacteria (2.5 μg mL^−1^ chlorophyll) resuspended in fresh medium sparged with 2% CO_2_ in air were dark-adapted for 3 min before measuring. The apparent quantum yield of photosystem II (Φ_II_), (F’_M_ - F_S_)/(F’_M_), and the coefficient of photochemical quenching (q_p_), (F’_M_ - F_S_)/(F’_M_ - F’_0_), were measured using a 1.5 s saturating pulses of actinic light (∼5,000 μmol photons m^−2^ s^−1^).

### 2.6. PSI absorbance changes

To evaluate P700 redox changes, samples were monitored semi-simultaneously with fluorescence measurements, using the instrument described in (Hall et al., 2013) by measuring absorbance changes at about 703 nm. The measuring beam was generated by a pulsed LED (720 nm peak emission, Rebel LUXEON Far Red) filtered with a 5 nm band pass filter centered at 700 nm, resulting in a measured emission peak at approximately 703 nm (Abramson et al., 2016). The signals were detected with a photodiode filtered with a Schott RG-695 filter to block actinic light (Hall et al., 2013). Samples containing cyanobacteria cells were prepared by resuspended cells in fresh medium to a concentration of 5 μg mL^−1^ chlorophyll and sparged with 2% CO_2_ in air. The samples were dark-adapted for 3 min before starting measurements. The percentage of oxidized P700 was calculated as [(P_ox_ - P_ss_)/(P_red_ - P_ox_)] · 100, where P_ox_ was the maximum extent of P700 absorbance signal induced by a saturating pulse of light during ∼0.5 s (∼5,000 μmol photons m^−2^ s^−1^); P_ss_, taken to be the fraction of P700^+^ in under steady state illumination, estimated by the extent of P700 absorbance signal induced by a short dark interval, and assuming that P700 reaches full reduction; and P_red_ is the level of P700 reduced in the dark. PSI traces were normalized to the last point of the steady-state level of oxidation (P_ss_).

### 2.7. Dark-interval relaxation kinetics (DIRK) absorbance changes

Steady-state levels of photooxidized P700 (P700^+^) were estimated by DIRK analysis (Sacksteder and Kramer, 2000), after 21 s of actinic illumination at three different intensities (100, 275 and 500 μmol photons m^−2^ s^−1^). Samples containing cyanobacteria (2.5 μg mL^−1^ chlorophyll) resuspended in fresh medium containing 2% CO_2_ and were dark-adapted for 3 min before measuring. The half-time of P700^+^ re-reduction (Tau), was measured from the absorbance change at 703 nm (ΔA_703_) during a dark interval of 2100 ms. Tau was calculated by the monotonic decay kinetic of ΔA_703_ produced after extinguishing the actinic light (Baker et al., 2007).

### 2.8. Microscopy and Image Analysis

All live-cell microscopy was performed on cells in exponential growth by centrifuging 2 mL of culture at 10,000 × *g* for 5 min, resuspending into 80 μL of BG11, and transferring a 2 μL aliquot to a 3% agarose pad. The cells were allowed to briefly equilibrate and be absorbed by the agarose (≥10 min) before the pad was placed onto a #1.5 glass coverslip for imaging. Images were captured using a Zeiss Axio Observer D1 inverted microscope equipped with an Axiocam 503 mono camera and a Zeiss Plan Apochromat 63X 1.4 NA oil-immersion lens.

Image analysis was done in OMERO (Allan et al., 2012) and Python 3. Cell segmentation was conducted using Cellpose (Stringer et al., 2021). Mean fluorescence intensity plots were generated as previously described (Sakkos et al., 2021). Briefly, cells were segmented using the chlorophyll autofluorescence channel, rotated such that the medial axis was horizontal, rescaled to ensure consistent boundaries, and the RbcS-mNG pixel intensity was averaged from each cell in the collection of images from its respective induction condition and time point. Foci locations were determined with a peak-finding algorithm using the Python package Photutils (Bradley et al., 2016).

### 2.9. Cellular viability

Cellular death was quantified using SYTOX Blue (1 mM in DMSO; Invitrogen, S34857) on a flow cytometer or SYTOX Orange (250 μM in DMSO; Invitrogen, S34861) and visualized by microscopy. SYTOX dyes are nucleic-acid-specific stains that are unable to reach the intracellular space due to the intact cell membrane of a non-damaged cell. At each time-point, 1 μL of the SYTOX blue working stock was added to 1 mL of culture (final concentration of 1 μM for SYTOX Blue or 250 nM for SYTOX Orange) and incubated in the dark at room temperature for 15 min. Heat-treated cyanobacterial cells were used as a positive control for dead cells. After incubation, 200 μL aliquots of the cultures were transferred to a 96-well plate to measure viability in the flow cytometer. Samples were collected on a 4-laser Attune CytPix with a CytKick Max Autosampler Software 6.2.0. The following optical configuration was used for each fluorophore (excitation|emission): SYTOX Blue [405nm|450/40] (BL1-A, 305V), Chlorophyll [405nm|660/20] (486V). Cyanobacterial samples were gated using FSC (400V) and SSC (250V) to distinguish the singlet population and with the chlorophyll to remove debris and noise, more than 10,000 cells were measured per sample type unless stated otherwise. Gating regions representing both intact, SYTOX-negative and membrane-damaged, SYTOX-positive cells were created in two-dimensional dot-plots (forward side scatter versus blue fluorescence). The raw data were analyzed using the FCS Express 7 software (De Novo Software, USA). For the microscope images, samples were processed following a protocol described in section 2.8.

### 2.10. Membrane inlet mass spectrometry (MIMS)

Online measurements of gas exchange were monitored using a mass spectrometer (model Prima PRO, Thermo Scientific). The membrane inlet system, consisting of a modified DW1 oxygen electrode chamber (Hansatech Instruments Ltd.) water-jacketed thermoregulated at 30 °C, was attached to the vacuum line of a mass spectrometer through a thin gas-permeable PTFE membrane (0.0125 mm) sealing the bottom of the chamber. ^18^O_2_ (isotope purity >98%; CK Gas Products Ltd.) tracing was used to discriminate O_2_ uptake and O_2_ production by PSII. ^16^O_2_ (*m/z* 32), ^18^O_2_ (*m/z* 36) and CO_2_ (*m/z* 44) were recorded with a time resolution of around 4 s. Samples were evenly mixed by constant stirring using a cross-shaped magnetic stirrer. A 2 mL aliquot of a cell suspension (10 μg mL^−1^ Chl*a*) was placed in the measuring chamber and prior to the measurement, cells were supplemented with ^18^O_2_ at an equivalent concentration to ^16^O_2_ and with 1.5 mM NaHCO_3_. Then, samples were measured for 5 min in darkness to record oxygen consumption caused by respiration. Following this period, actinic light (500 μmol photons m^−2^ s^−1^) was applied via a 150-W, 21-V EKE quartz halogen-powered fiber optic illuminator (Fiber-Lite DC-950; Dolan-Jenner). Gas-exchange kinetics and rates were determined according to (Beckmann et al., 2009). Final Chl*a* concentration, determined spectrophotometrically in 100% methanol according to Porra et al. (Porra et al., 1989), was conducted at completion of each measurement for standardizing the calculated gas exchange rates.

### 2.11. Determination of intracellular glycogen content

Glycogen content was determined as described previously with minor modifications (Gründel et al., 2012). Cyanobacterial culture aliquots (2 mL) were pelleted down by centrifuging at 5,000 *xg* for 10 minutes. Pellets were flash-frozen in liquid nitrogen and were stored at -80 °C until extraction. For isolation of glycogen, the pellets were resuspended in 200 μL 30% (w/v) KOH and incubated in a heat block at 95 °C for 2 hours. Samples were cooled down on ice. Complete precipitation of glycogen was achieved by the addition of 600 µL of cold absolute ethanol and overnight incubation at −20 °C. The precipitated glycogen was recovered by centrifugation at 17,000 *xg* for 15 min at 4 °C. The supernatant was removed, and the glycogen pellets were dried for 40 min at 60°C using a SpeedVac. The precipitated glycogen was resuspended in 200 µL of miliQ H_2_O by vortexing. The homogeneous samples were quantified using the EnzyChrome glycogen assay kit (BioAssay Systems, E2GN-100) according to the manufacturer’s instructions.

### 2.12. Transmission Electron Microscopy

At each time point, 2 ml samples were prepared by diluting cultures to a final OD_750_ 1.5. Cells were pelleted and fixed overnight at 4 °C with 2% glutaraldehyde/2% paraformaldehyde in phosphate buffer (pH 7.4), suspended into a 2% agarose bead and cut into ∼1 mm cubes. Following three washes with 0.1 M sodium cacodylate buffer, cells were suspended in 1% osmium tetroxide/1.5% potassium ferrocyanide and incubated overnight at 4 °C. After incubation, cells were washed with HPLC-quality H_2_O until they appear clear. Cells were then suspended in 1% uranyl acetate and microwaved for 2 min using a MS-9000 Laboratory Microwave Oven (Electron Microscopy Science), decanted, and washed until clear. Cells were dehydrated in increasing acetone series (microwave 2 min) and then embedded in Spurr’s resin (25% increments for 10 min each at 25 °C). A final overnight incubation at room temperature in Spurr’s resin was done, then cells were embedded in blocks which were polymerized by incubation at 60 °C for 3 days. Thin sections of approximately 50 nm were obtained using an MYX ultramicrotome (RMC Products), post-stained with 1% uranyl acetate and Reynolds lead citrate, and visualized on a JEM 100CX II transmission electron microscope (JEOL) equipped with an Orius SC200-830 CCD camera (Gatan).

### 2.13. Quantification of Reactive Oxygen Species (ROS)

ROS were quantified by using the fluorescent marker H_2_DCFDA (2′,7′-dichlorodihydrofluorescein diacetate; Invitrogen, D399) as previously reported (Diamond et al., 2017). Briefly, 2 mL of the cultures were collected and split into 1-mL aliquots. H_2_DCFDA was added to one sample at a final concentration of 5 μM. Tubes were protected from light and shaken at 30 °C for 30 min. After incubation, 200 μL of each tube was added to a separate well in a 96-well plate. The fluorescent product 2’,7’-dichlorofluorescein (DCF) was monitored via microplate reader (excitation 480 nm, emission 520 nm; SpectraMax M2 microplate reader by Molecular Devices). Untreated-sample background fluorescence was then subtracted from treated-sample fluorescence values and fluorescence data were normalized to OD_750_ of each sample.

### 2.14. Determination of growth of cyanobacterial cultures in the presence of hydrogen peroxide

To determine the growth of the different cyanobacterial strains in the presence of H_2_O_2_, cell cultures were adjusted to an OD_750_ of 0.3. Hydrogen peroxide, at final concentrations ranging from 50 μM to 2 mM, was added to 1.5-mL aliquots of cultures in separate wells of a 24-well plate or to 50-mL flasks with cultures. Images of the plates, as well as fluorescence microscopy images, were taken after a 24-h incubation under standard growth conditions.

### 2.15. Statistical analysis

Recorded measurements are represented as mean values, with error bars expressing the SD of n≥3 biological replicates experiments, as indicated. The significance of differences between groups was evaluated by one-way ANOVA followed by Tukey’s multiple comparison test or by an unpaired Student’s t-test. Statistical analyses were carried out using GraphPad Prism software (GraphPad Software Inc., San Diego, CA). Differences were considered statistically significant at P < 0.05.

## 3. Results

### 3.1. Remodeling carbon fixation machinery following sucrose secretion requires *rpaA*

As discussed in the introduction, we recently implicated RpaA as a two-component protein that appears to be involved in the regulation of photosynthetic changes following activation of a heterologous sucrose secretion pathway (Santos-Merino et al., 2024). To further investigate the potential roles of RpaA in adapting photosynthesis and the CCM we first validated the impact of loss-of-function mutants (Δ*rpaA*) in a strain expressing a carboxysome reporter (RbcS-mNG) and bearing an inducible sucrose-export pathway (CscB-SPS^export^). We confirmed that Δ*rpaA* strains lacked the characteristic rise in photosynthetic activity following activation of sucrose secretion, as measured by apparent quantum efficiency of photosystem II (Φ_II_; Figure 1A). Similarly, deletion of *rpaA* abrogated the increase in carboxysome number and size that follows sucrose secretion (Singh et al., 2022), as monitored by the carboxysome reporter strain (Figures 1B an 1C and Supplementary Figure S1). Importantly, the Δ*rpaA* background growth rate and other photosynthetic parameters appeared comparable to WT under the laboratory cultivation conditions with constant light, although Δ*rpaA* cell size was slightly diminished (Supplementary Figure S2). These results are consistent with other recent reports that have demonstrated that Δ*rpaA* strains perform similarly to wildtype under conditions of constant light and high CO_2_; sometimes even exceeding wildtype growth (Espinosa et al., 2015; Diamond et al., 2017; Puszynska and O’Shea, 2017). Therefore, deletion of *rpaA* eliminates the enhancements in photosynthesis and increased carboxysome number typically observed in cells induced to export sucrose (Figures 1 and Supplementary Figure S1) but does not globally misregulate essential pathways required for growth and division under controlled laboratory conditions (Supplementary Figure S2).

**Figure 1.**
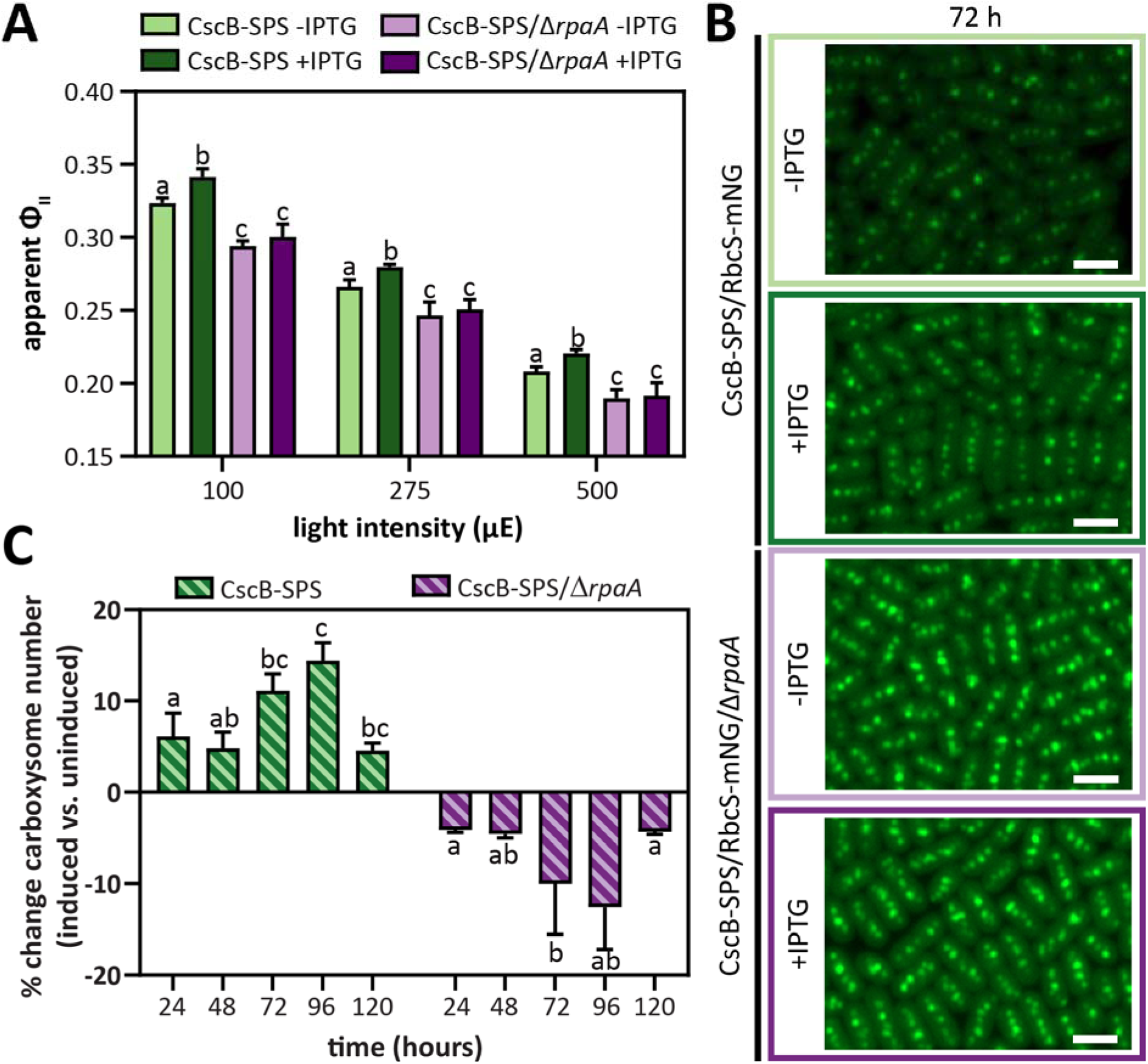
Acclimatory changes in the photosynthetic machinery following sucrose export are absent in Δ*rpaA*. **(A)** Apparent Φ_II_ values measured at three different light intensities 24 hours after the induction of sucrose export in the strain CscB-SPS^export^ in the presence/absence of RpaA. **(B)** Carboxysome number change (expressed as percentage) after induction of sucrose export in the strain CscB-SPS^export^ in the presence/absence of RpaA. **(C)** Carboxysomes at 72 hours visualized via the RbcS-mNG reporter in the strain CscB-SPS^export^ in the presence/absence of RpaA. Scale bar: 2 µm. **(A, B)** Averages of ≥3 independent biological replicates are shown + SD. Significance was calculated by one-way ANOVA followed by Tukey’s multiple comparison test. Data points labeled with different letters are significantly different (P < 0.05).

### 3.2. Δ*rpaA* mutants exhibit growth arrest and dramatic carboxysome disassembly under mixotrophic conditions

As we and others have shown that carboxysome structure and composition is impacted by growth under mixotrophic conditions (Muth-Pawlak et al., 2022; Singh et al., 2022), we next examined the impact of *rpaA* knockout on carboxysome restructuring under mixotrophy. We encoded the sucrose permease gene under the IPTG-inducible P*_trc_* promoter to allow genetic control over sucrose import (CscB^import^) and supplemented the growth medium with sucrose. In a wildtype background, mixotrophic growth mildly suppressed carboxysome number and increased the incidence of carboxysome clustering (Figure 2A), consistent with prior studies (Singh et al., 2022). By contrast, under conditions where CscB was activated, allowing uptake of extracellular sucrose, Δ*rpaA* strains exhibited an immediate growth arrest and carboxysome reorganization followed by a dramatic disassembly of carboxysomes within 1-3 days of the onset of mixotrophic growth (Figures 2A-2C and Supplementary Figure S3). The loss of carboxysomal integrity appeared to be gradual, with some Δ*rpaA* cells displaying an increase in delocalized Rubisco signal throughout the cytosol in the population as early as 24 hours, but only a fraction of cells exhibiting a complete loss of carboxysomal puncta (Supplementary Figure S3A). The fraction of cells with diminished carboxysome puncta and increased cytosolic Rubisco localization increased over time, with nearly all cells lacking carboxysomes after 72 hours of sucrose feeding (Figure 2A). Finally, we observed that cells displaying increased autofluorescence of natural cyanobacterial pigments (possibly associated with impaired photosynthetic performance) were associated with a more rapid loss of carboxysome puncta over time (Figure 2A).

**Figure 2.**
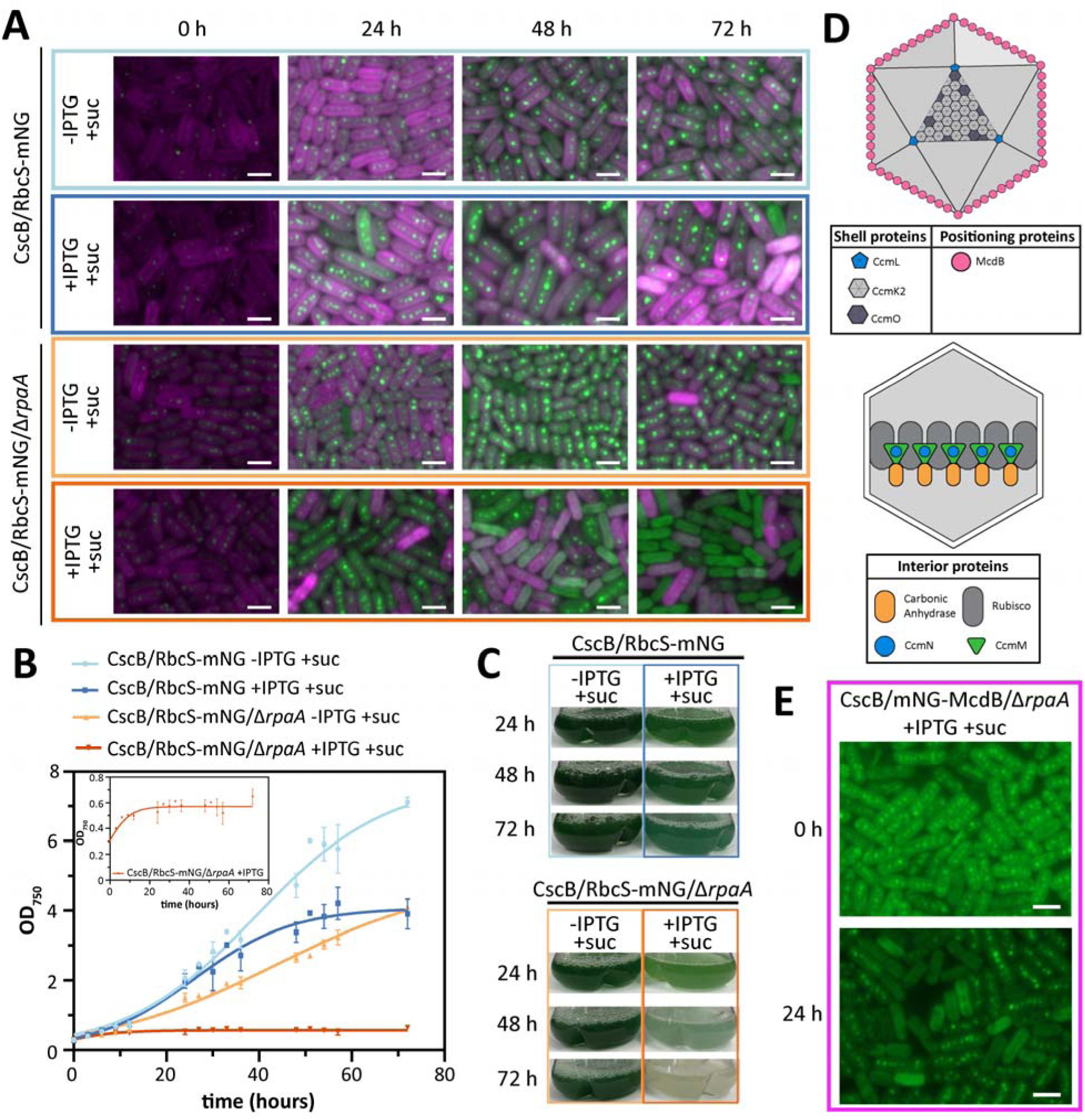
Mixotrophic conditions induces carboxysome breakdown and growth arrest in Δ*rpaA*. **(A)** Time-course of carboxysome status under photoautotrophic and mixotrophic conditions in the strain CscB^import^ in the presence/absence of RpaA. Scale bar: 2 µm. **(B)** Growth curves of the strain CscB^import^ in the presence/absence of RpaA in photoautotrophic and mixotrophic conditions with zoom in of the growth curve of CscB/RbcS-mNG/Δ*rpaA*. Averages of ≥3 independent biological replicates are shown ± SD. **(C)** Appearance of the cultures of the strain CscB^import^ in the presence/absence of RpaA under photoautotrophic and mixotrophic conditions. **(D)** Cartoon illustration of internal and external carboxysome components. **(E)** Carboxysome status under mixotrophic conditions in the strain CscB^import^ in the presence/absence of RpaA by tracking mNG-McdB. Scale bar: 2 µm.

The observed loss of puncta following mixotrophic growth conditions must be attributed to dissolution of existing carboxysomes rather than failure to build new microcompartments because of the complete growth arrest observed (Figure 2B). To better assess the dynamics of carboxysome breakdown, we monitored dynamics of an endogenously tagged outer carboxysome component maintenance of carboxysome distribution B (McdB)(Figure 2D), a ParB-family protein involved in microcompartment positioning (MacCready et al., 2018). McdB utilizes a conserved motif to bind to the cytosolic-facing side of bacterial microcompartment shell proteins, and it is therefore a relatively peripheral component of the carboxysome (Basalla et al., 2024). Within 24 hours following onset of mixotrophic growth in Δ*rpaA* cells, we observed a near complete loss of McdB-mNG with the carboxysome shell in nearly all cells (Figure 2E). By comparison, the Rubisco core of carboxysomes remained present at this time point in most cells (Supplementary Figure S4), suggesting that carboxysome disassembly may initiate with components associated with the shell proteins and proceed inward. Consistent with the loss of the positional machinery, we observed that Rubisco puncta in mixotrophic Δ*rpaA* cells were frequently mispositioned relative to one another or localized to the cell pole (Supplementary Figure S5). The localization of Rubisco to cell poles has been proposed to be an initiating step of carboxysome disassembly in prior studies (Hill et al., 2020).

### 3.3. Δ*rpaA* cells undergo a severe photosynthetic impairment under mixotrophy

The physiological characteristics of *rpaA* mutants grown under mixotrophic conditions (*i.e.*, cell growth arrest, bleaching of pigments, and carboxysome breakdown) (Figure 2 and Supplementary Figures S3 and S4) suggest that photosynthetic processes may be misregulated (Sunil et al., 2013; Calzadilla and Kirilovsky, 2020). We directly assessed photosynthetic performance of cultures grown under photoautotrophic and mixotrophic conditions via fluorimetry. In sucrose-fed Δ*rpaA* lines, the openness of PSII reaction centers (q_p_) was dramatically decreased at all tested levels of illumination relative to modest changes in sucrose-fed control strain CscB/RbcS-mNG (Figure 3A). Similarly, the apparent Φ_II_ was reduced almost to zero under mixotrophic growth in Δ*rpaA* cultures, although Φ_II_ was also partially suppressed by sucrose feeding in the reference control (Figure 3B). Other measured photosynthetic values were not as strongly impacted by sucrose feeding in either line, including the oxidation state of PSI or the estimates of electron flux from cytochrome *b_6_f* (Supplementary Figures S6A and S6B), overall indicating that PSII and/or electron flux near PQ were the most strongly impacted.

**Figure 3.**
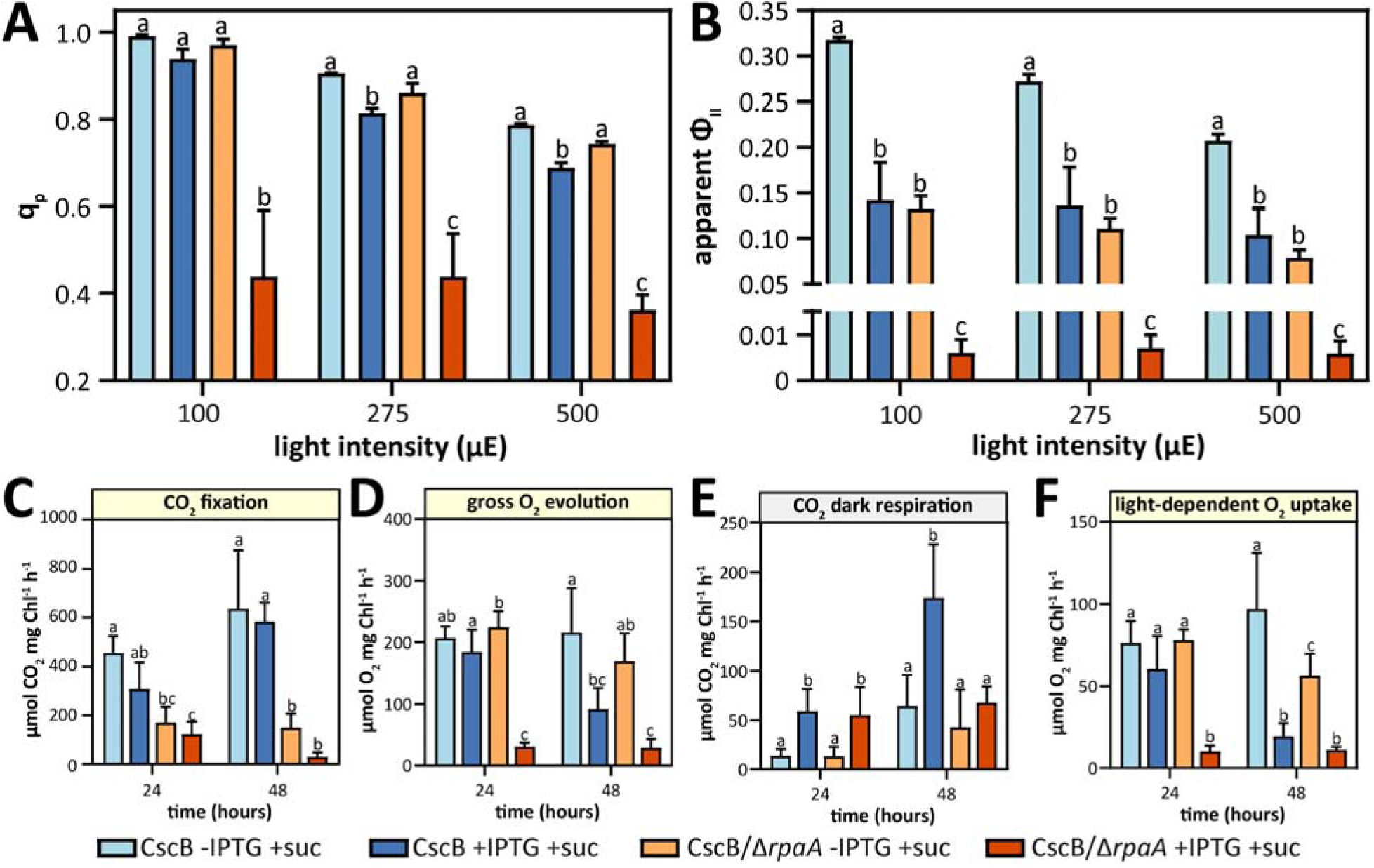
Mixotrophic growth conditions impair PSII activity and have different effects on the rate of O_2_ and CO_2_ fluxes. **(A)** q_p_ values. **(B)** Apparent Φ_II_ values. Quantification of the steady state fluxes rates shown in Supplementary Figures S6C-S6H for **(C)** CO_2_ fixation, **(D)** gross O_2_ evolution, **(E)** CO_2_ dak respiration, and **(F)** light-dependent O_2_ uptake for the strains CscB/RbcS-mNG and CscB/RbcS-mNG/Δ*rpaA*. CO_2_ exchange rates in the strains CscB/RbcS-mNG and CscB/RbcS-mNG/Δ*rpaA* under photoautotrophic and mixotrophic conditions. Averages of ≥3 independent biological replicates are shown + SD. Significance was calculated by one-way ANOVA followed by Tukey’s multiple comparison test. Data points labeled with different letters are significantly different (*P* < 0.05).

To gain additional insight into photosynthetic fluxes in control and Δ*rpaA* lines, we turned to membrane inlet mass spectrometry (MIMS), which can disentangle CO_2_ and O_2_ gas exchange rates originating from photosynthetic and respiratory metabolism. Consistent with the sharp decline in q_P_ and Φ_II_, we observed a rapid loss of photosynthetic CO_2_ fixation under both photoautotrophic and mixotrophic growth conditions in Δ*rpaA* cells (Figure 3C and Supplementary Figures S6C and S6D), accompanied by a loss of O_2_ evolution only under mixotrophic conditions in the Δ*rpaA* strain (Figure 3D and Supplementary Figures S6E and S6F). Continued CO_2_ respiration was observed in the Δ*rpaA* cells under dark conditions (Figure 3E and Supplementary Figures S6C and S6D), providing direct evidence that the cells remain metabolically active during growth arrest (Figure 2). Almost a complete loss of O_2_ uptake induced by light was observed under mixotrophic conditions (Figure 3E and Supplementary Figures S6G and S6H), suggesting that reducing equivalents accumulated on the pETC are not efficiently consumed by alternative electron pathways that normally quench excess reductant by donating them to molecular oxygen (*i.e.*, water-water cycles) in the Δ*rpaA* line (Nikkanen et al., 2021).

Given the severity of the photosynthetic impairment that mixotrophic conditions initiated in the Δ*rpaA* line, we used the vital dye SYTOX to confirm that cells exhibiting carboxysome breakdown remained viable. Fluorescence-activated cell sorting (FACS) showed that cell viability began to be impaired 48 hours under mixotrophic growth, with no viable cells after 72 hours (Figure 4A and Supplementary Figure S7). This indicated that initiation of carboxysome disassembly (*i.e.*, evident at 24 hours) precedes that of the loss of cell viability (between 48 and 72 hours under mixotrophic growth). Fluorescence microscopy with SYTOX confirmed this interpretation, as cells that show disruption of carboxysomes remain SYTOX negative 24 and 48 hours into the mixotrophic-induced growth arrest (Figure 4B and Supplementary Figure S5).To further evaluate the interplay between carboxysome breakdown and cell viability in Δ*rpaA* cells, we fed sucrose for 60 hours (when nearly all Δ*rpaA* cells have no evident carboxysomes), then tracked recovery by returning cells to photoautotrophic growth (Supplementary Figure S8). Following sucrose removal, Δ*rpaA* cells exited the growth arrest (<24 hours), recovered pigmentation (24-48 hours), and reassembled carboxysomes (24-72 hours) (Supplementary Figures S8A-S8E). Whereas growth of Δ*rpaA* cells immediately resumed when sucrose was removed (Supplementary Figure S8D), recovery of cellular pigmentation and carboxysome puncta was delayed (Supplementary Figures S8A-S8C). A further indication that growth arrested cells remained metabolically active was the greatly-elevated ROS levels in sucrose-fed Δ*rpaA* cultures; ROS levels gradually returned to the baseline of paired controls by 72 hours following removal of sucrose (Supplementary Figure S8E).

**Figure 4.**
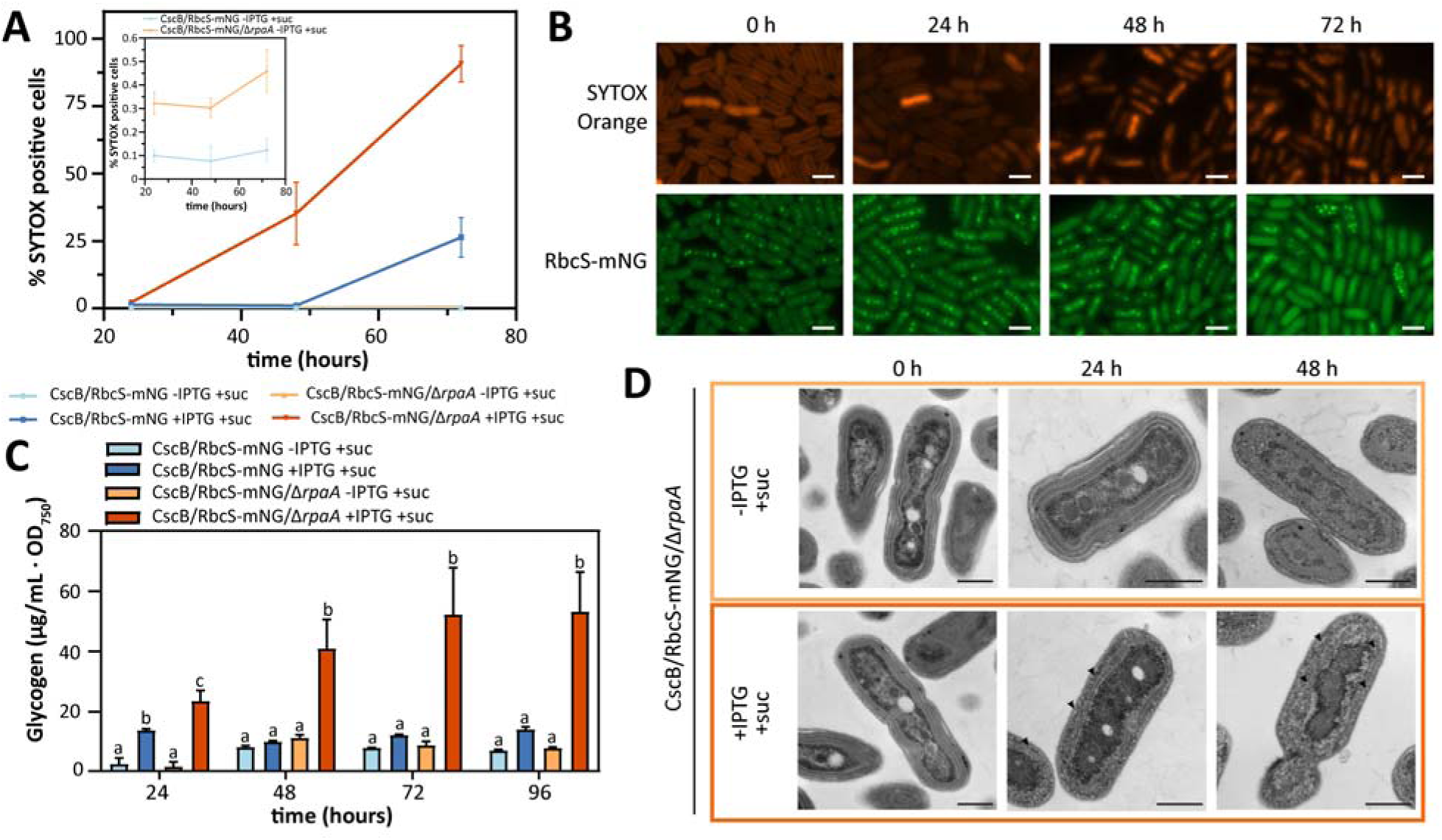
Carboxysome disassemble precedes cell death and is accompanied by glycogen accumulation. **(A)** Percentage of SYTOX Blue-positive cells (dead cells) measured by flow cytometry under photoautotrophic and mixotrophic conditions, with detail of CscB/RbcS-mNG and CscB/RbcS-mNG/Δ*rpaA* under photoautotrophic conditions. Averages of ≥3 independent biological replicates are shown ± SD. **(B)** Time-course of SYTOX Orange-positive cells and carboxysomes by tracking RbcS-mNG under mixotrophic conditions in the strain CscB^import^ in the absence of RpaA. Scale bar: 2 µm. **(C)** Quantification of glycogen per OD_750_ unit at different time points under photoautotrophic and mixotrophic conditions in the CscB strain in the presence and absence of RpaA. Averages of ≥3 independent biological replicates are shown + SD. Significance was calculated by one-way ANOVA followed by Tukey’s multiple comparison test. Data points labeled with different letters are significantly different (*P* < 0.05). **(D)** Transmission electron microscopy of cells of the strain CscB^import^ in the absence of RpaA grown photoautotrophically and mixotrophically; black arrows indicate glycogen granules. Scale bar: 500 nm.

RpaA has recently been implicated in regulation of central carbon metabolism and glycogen mobilization (Diamond et al., 2015; Diamond et al., 2017; Puszynska and O’Shea, 2017; Santos-Merino et al., 2024), so we examined the utilization of sucrose in Δ*rpaA* lines. We observed that sucrose feeding leads to glycogen accumulation in both wild-type (WT) and Δ*rpaA* backgrounds in the first 24 hours. However, while WT background restored normal glycogen levels by 48 hours, Δ*rpaA* lines exhibited a continual accumulation of glycogen over time (Figure 4C). Indeed, glycogen hyperaccumulation in sucrose-fed Δ*rpaA* cells was directly observable by electron microscopy, which revealed that glycogen bodies proliferated throughout the cytosol and within the thylakoid membranes (Figure 4D). Taken together, fluorescence kinetics, MIMS, vital dyes, and electron microscopy suggest that mixotrophic growth in Δ*rpaA* line induces severe impairment of photosynthetic reactions, arrests cell growth while maintaining metabolic activity, triggers abnormal levels of glycogen deposition, initiates carboxysome disassembly, and eventually can lead to cell death under prolonged exposure.

### 3.4. PQ overreduction leads to H_2_O_2_ formation and carboxysome breakdown

A variety of stressors can lead to imbalances within the pETC, often leading to activation of photoprotective mechanisms including alternative pathways to quench excess reductant to avoid formation of ROS and photodamage (Pospisil, 2016). In the Δ*rpaA* mutant, we observe several signs that indicate an accumulation of electrons in the pETC, including a drop in the values of q_p_ and in the apparent Φ_II_ (Figures 3A and 3B), and the loss of gross O_2_ evolution (Figures 3D and 3F and Supplementary Figures S6E-S6H). Moreover, the suppression of light-induced O_2_ uptake we observed in mixotrophic Δ*rpaA* strains (Figure 3F) suggests that alternative pathways for quenching electrons in the pETC are suppressed (*e.g.*, flavodiiron proteins Flv1/3), which could further exacerbate the buildup of reductant (Diamond et al., 2017).

Because of the indicators of potential imbalances in the pETC, we directly monitored ROS production in Δ*rpaA* cells under mixotrophic conditions and observed a sharp increase in ROS production that peaked at 72 hours after initiation of sucrose feeding (Figure 5A). To further dissect the phenotype, we used chemical inhibitors and growth conditions to modulate redox status of the pETC. Treatment of Δ*rpaA* cells with 3-(3,4-dichlorophenyl)-1,1-dimethylurea (DCMU), a specific inhibitor of PSII that blocks electron transport from Q_A_ to Q_B_ and thereby oxidizes the pETC at PQ and downstream electron carriers, decreased ROS production under mixotrophic conditions and prevented carboxysome breakdown (Figures 5A and 5C and Supplementary Figure S10C). In parallel, sucrose feeding of Δ*rpaA* in the dark, where photosynthesis is inactive (Khorobrykh et al., 2020), also prevented both ROS production and preserved carboxysome integrity. By contrast, treatment with dibromothymoquinone (DBMIB) blocks electron transfer from PQ to Cytochrome *b_6_f,* did not rescue sucrose-fed Δ*rpaA* cells, which continued to strongly produce ROS and exhibit carboxysome breakdown (Figures 5A and 5C and Supplementary Figure S10C). Darkness and DCMU treatment also recovered the loss in cell viability and pigmentation usually observed in mixotrophic Δ*rpaA* cells (Supplementary Figure S10). Taken together, our results suggest overreduction of PQ in Δ*rpaA* cultures is associated with ROS formation and carboxysome breakdown.

**Figure 5.**
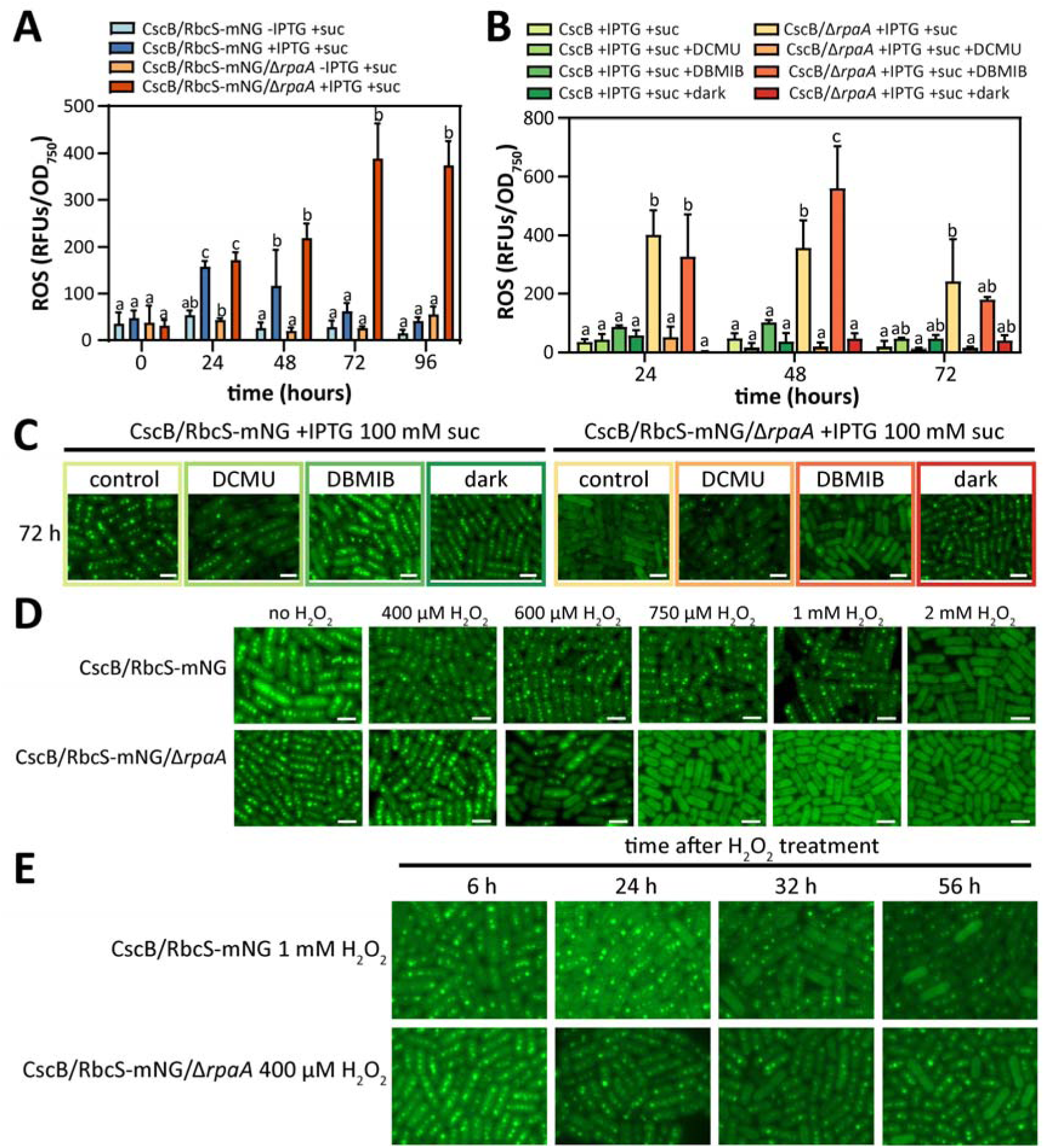
Reduction of the PQ pool leads to accumulation of ROS and carboxysome disassembly. Quantification of cellular ROS accumulation measured by H_2_DCFDA fluorescence at different time points **(A)** following activation of sucrose import, and **(B)** following the addition of photosynthesis inhibitors or growing the cells in darkness in sucrose feeding conditions in the CscB^import^ strain in the absence of RpaA. **(C)** Carboxysome status at 72 hours in response to photosynthesis inhibitors or growing the cells in darkness in sucrose feeding conditions in the CscB^import^ strain in the presence/absence of RpaA by tracking RbcS-mNG in the fluorescence microscope. Scale bar: 2 µm. **(D)** Carboxysome status after 24 hours exposure to different concentrations of H_2_O_2_ in the CscB^import^ strain in the presence/absence of RpaA by tracking RbcS-mNG. Scale bar: 2 µm. **(E)** Carboxysome status after exposure to 1 mM of H_2_O_2_ for CscB^import^ strain and 400 µM H_2_O_2_ for CscB^import^ strain in the absence of RpaA by tracking RbcS-mNG. Scale bar: 2 µm. **(A, B)** Averages of ≥3 independent biological replicates are shown + SD. Significance was calculated by one-way ANOVA followed by Tukey’s multiple comparison test. Data points labeled with different letters are significantly different (*P* < 0.05).

Hydrogen peroxide (H_2_O_2_) is a dominant form of ROS that has established signaling roles in plants (Mubarakshina and Ivanov, 2010), although any conserved regulatory function in cyanobacteria has not been clearly established (Latifi et al., 2009). We therefore tested direct application of H_2_O_2_, observing that this treatment also impacted cellular pigmentation and carboxysome integrity in both WT and Δ*rpaA* strains (Figure 5D). Interestingly, the Δ*rpaA* mutant exhibited increased sensitivity to external addition of H_2_O_2_ in terms of cell viability, pigmentation, and carboxysome integrity (Figure 5D and Supplementary Figure S11). Acute treatment with H_2_O_2_ (2 mM H_2_O_2_ for WT or 750 µM H_2_O_2_ for Δ*rpaA*) led to a rapid bleaching and disassembly of most carboxysomes within the first 24 hours. After acute H_2_O_2_ treatment, those cells that still contained visible RbcS-mNG puncta typically possessed only one or two carboxysomes which were typically located in the pole of the cells (Supplementary Figure S11D); this localization pattern has been previously reported as preceding carboxysome degradation (Hill et al., 2020). Yet, cell cultures did not always recover growth following acute H_2_O_2_ treatment, complicating interpretation of these results. We therefore conducted further analysis using H_2_O_2_ treatments at concentrations where cell growth and pigmentation were unaffected (Supplementary Figure S11A). Cells treated with subacute H_2_O_2_ (1 mM for WT or 400µM for Δ*rpaA*) did not exhibit substantive changes in growth or pigmentation (Supplementary Figure 12). However, in the hours immediately following subacute H_2_O_2_ treatment, subtle alterations in carboxysome number and RbcS-mNG puncta organization were observed (Figure 5E). Crucially, at later time points (36-72 hours post H_2_O_2_ exposure), carboxysome phenotypes appeared exacerbated, with many cells in the population displaying heterogeneity in carboxysome brightness, mispositioned/polar carboxysome puncta, too few carboxysomes or no carboxysome puncta at all (Figure 5E and Supplementary Figure S12). The carboxysome phenotypes at later time points (≥ 36 hours) following subacute H_2_O_2_ treatment was less severe but were similar in character and timing to phenotypes observed under mixotrophic growth (>24 hours).

### 3.5. Physiological conditions that lead to ROS generation are associated with carboxysome rearrangement

We next asked if other physiologically-relevant environmental conditions are associated with carboxysome disassembly beyond mixotrophic growth and we monitored any correlation with redox imbalance and ROS production. Light and inorganic carbon availability are two critical environmental factors strongly influencing cyanobacterial photosynthetic performance and ROS generation (Khorobrykh et al., 2020; Krieger-Liszkay and Shimakawa, 2022), therefore we revisited WT and Δ*rpaA* cells under ambient CO_2_ conditions while varying both light intensity (HL, 150 μmol photons m^−2^ s^−1^; LL, 17 μmol photons m^−2^ s^−1^) and sucrose feeding. At ambient CO_2_ levels and HL, we found that Δ*rpaA* displayed substantial carboxysome breakdown even in the absence of sucrose feeding (Figure 6A, HL). Ambient CO_2_ alone was insufficient to induce carboxysome disassembly in Δ*rpaA* cells, as carboxysomes were well maintained in LL conditions both under photoautotrophic and mixotrophic conditions (Figure 6A, LL). We also detected an increase in ROS accumulation that was more pronounced under photoautotrophic conditions at both HL and LL in comparison to mixotrophic conditions (Figure 6B), though the absolute values of detected ROS were substantially higher across all tested cultures under air relative to 3% CO_2_ (Figure 5A). Intriguingly, mixotrophic growth partially rescued the loss of carboxysomes in Δ*rpaA* cells under ambient CO_2_ and HL (Figure 6A). In each of these cases, elevated ROS was strongly correlated with carboxysome disassembly (Figure 6B), as well as chlorophyll content and cell viability, although growth arrest was not observed under these conditions (Supplementary Figures S13 and S14).

**Figure 6.**
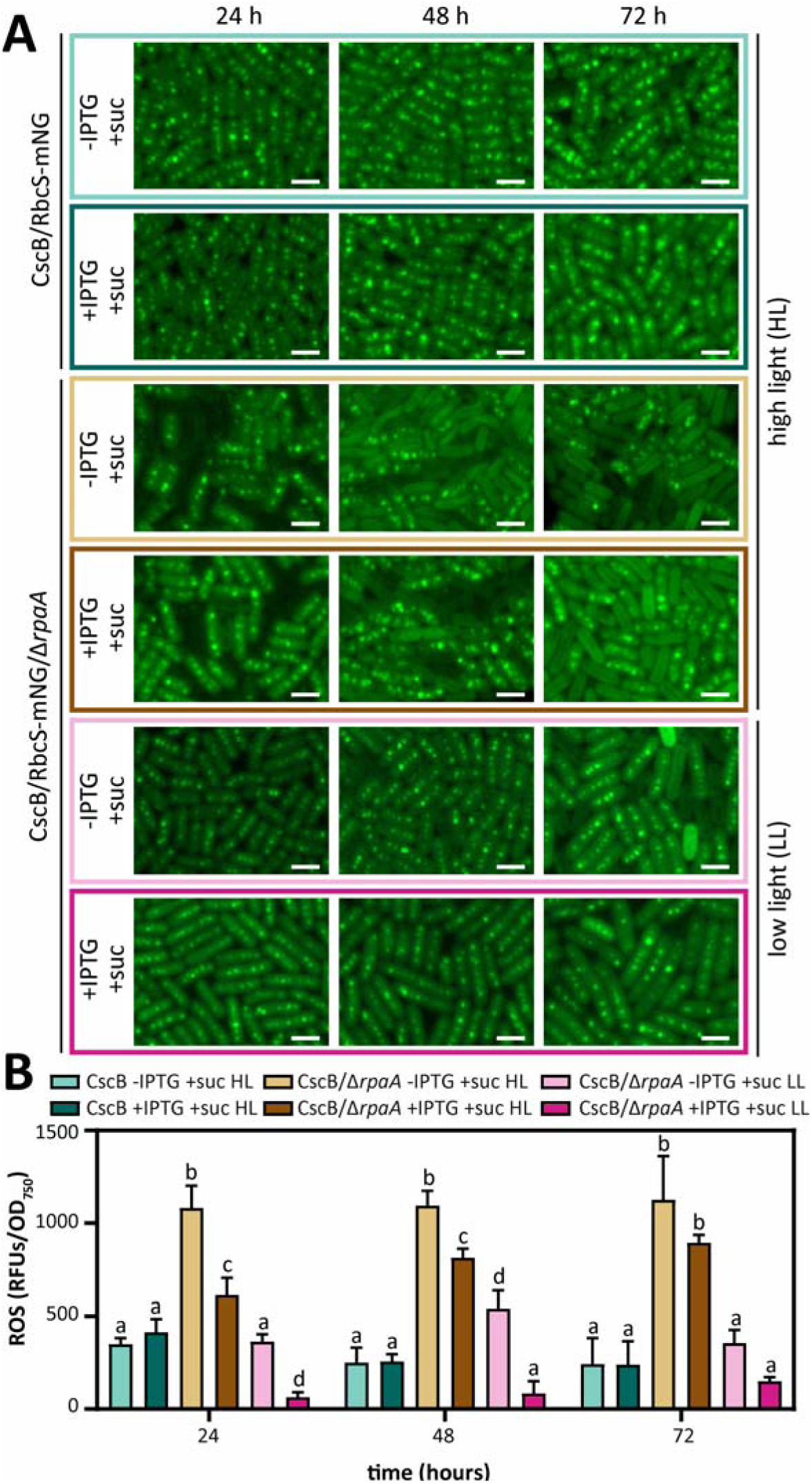
Carboxysome breakdown is delayed under ambient air conditions and low light intensity. **(A)** Carboxysome status in response to ambient air conditions under mixotrophic conditions for the strains CscB/RbcS-mNG and CscB/RbcS-mNG/Δ*rpaA* by tracking RbcS-mNG. Scale bar: 2 µm. **(B)** Quantification of cellular ROS accumulation measured by H_2_DCFDA fluorescence at different time points following the transference to ambient air conditions during sucrose feeding conditions for the strains CscB/RbcS-mNG and CscB/RbcS-mNG/Δ*rpaA*. Averages of ≥3 independent biological replicates are shown + SD. Significance was calculated by one-way ANOVA followed by Tukey’s multiple comparison test. Data points labeled with different letters are significantly different (*P* < 0.05).

## 4. Discussion

Taken together, our data supports a novel connectivity between the redox status of the PQ pool and carboxysome status is linked to RpaA function. Recently, we reported that deletion of RpaA prevented a number of acclimatory responses in the light reactions of photosynthesis (*e.g.*, increased O_2_ evolution and apparent Φ_II_) that are typically induced by activation of an engineered sucrose sink (Santos-Merino et al., 2024). Another photosynthetic impact we and others have observed following activation of sucrose export pathways is an increase in CO_2_ fixation rate that is associated with a change in carboxysome number and increased rubisco content (Ducat et al., 2012; Singh et al., 2022; Wang et al., 2023). Here, we show that Δ*rpaA* mutants also fail to reorganize carboxysomes following sucrose export, but instead can display dramatic reorganization and disassembly of carboxysomes under environmental conditions that lead to overreduction of the pETC and formation of ROS (Figures 2, 5 and 6 and Supplementary Figures S3, S4 and S12). Under prolonged conditions of H_2_O_2_ generation, carboxysome breakdown can be complete, although carboxysomes will reform in cells once the stress is removed (Figure 2 and Supplementary Figures S3 and S8). Importantly, chemical inhibitors that block photosynthetic reduction of the PQ pool prevent the carboxysome breakdown phenotype (Figure 5 and Supplementary Figure S10), suggesting that PQ redox status is especially important to levels of ROS and/or carboxysome integrity. Finally, we show that direct addition of H_2_O_2_ can initiate a reorganization or breakdown of carboxysomes *in vivo* (Figure 5D and Supplementary Figure S11).

One possible interpretation of our data is that the reduction status of PQ and associated ROS formation is part of a normal mechanism that cyanobacteria use to remodel the carboxysome in response to different environmental changes. It has now been well documented that the size, number, and subunit composition of carboxysomes is correlated with light and CO_2_ levels in a number of model cyanobacterial species (Sun et al., 2016; Rohnke et al., 2018; Sun et al., 2019; Rillema et al., 2021; Lucius and Hagemann, 2024). In one particularly relevant example, high light was shown to significantly increase in carboxysome number in *S. elongatus,* but carboxysome remodeling was partially blocked via addition of DCMU (Sun et al., 2016), although the mechanism for this block was unclear. It is also established that certain environmental conditions (*e.g.*, high light, low CO_2_), can lead to over-reduction of the PQ pool and resulting production of ROS (Lea-Smith et al., 2013). Our data extends upon prior publications by suggesting PQ-dependent carboxysome remodeling may be a function of ROS (potentially H_2_O_2_) derived from overreduction of this redox carrier.

Cyanobacteria usually exhibit constitutive expression and assembly of carboxysomes and (to our knowledge) the complete breakdown of carboxysomes is nor reported under physiological conditions elsewhere, although other studies have examined conditions where carboxysome integrity is compromised. First, live-cell imaging studies of cyanobacteria engineered to be gradually depleted of key structural components of the carboxysome have demonstrated that degrading carboxysomes are mislocalized from the nucleoid to the cell pole prior to the dissolution of the Rubisco core (Hill et al., 2020). This elegant imaging of the carboxysome lifecycle matches our observation of mispositioned and polar Rubisco puncta that precede carboxysome loss in Δ*rpaA* mutants (Figures 2, 5, and 6 and Supplementary Figures S3, S4, S5, S10, and S11). The cyanobacterial species, *Microcystis aeruginosa*, produces a toxin called microcystin that binds to Rubisco and interferes with its encapsulation within the carboxysome lumen, resulting in delocalized and thylakoid-associated Rubisco aggregates (Barchewitz et al., 2019). It has also been shown that H_2_O_2_ degrading activities of thioredoxin and peroxiredoxin are also inhibited by microcystin (Alexova et al., 2016; Schuurmans et al., 2018), which could sensitize *M. aeruginosa* to ROS-related pathways, including the carboxysome breakdown phenotype we report here.

Based on our data and prior literature Δ*rpaA* mutants appear to be particularly sensitized to redox imbalances and may thereby exhibit an amplified response to a redox related regulatory signal. RpaA is involved in controlling redox balance, as observed for the incapacity of Δ*rpaA* mutant to clear ROS out during the night, that have been excessively accumulated during the day (Diamond et al., 2017). The inability to detoxify ROS of the Δ*rpaA* cells makes them unable to survive the dark period, leading to a conditional light–dark lethality. As deletion of RpaA has been associated with misregulation of the expression of hundreds of genes in cyanobacteria (Markson et al., 2013), it is difficult to predict how directly RpaA is involved in mitigating or transducing ROS signals. However, Δ*rpaA* mutants are strongly downregulated in thioredoxin-dependent peroxidase (2-cys prx) (Markson et al., 2013; Puszynska and O’Shea, 2017), and Δ*rpaA* mutants exhibit a deficit in NADPH that is required for the activity of ROS scavenger enzymes (Diamond et al., 2017). Indeed, 2-cys prx is the main enzyme in *S. elongatus* responsible for detoxifying ROS and has been previously shown to be upregulated under photomixotrophic growth conditions (Perelman et al., 2003; Tan et al., 2022). More directly, we have recently shown enrichment of thioredoxin peroxidase and catalase in proximity labeling studies with RpaA (Santos-Merino et al., 2024), indicating a possible direct RpaA interaction that would require additional study. In this same interactome, we detected a putative RpaA-Flv3 interaction that may merit additional validation given that we observe a loss of light-induced O_2_ uptake catalyzed by the flavodiiron proteins Flv1/Flv3. Curiously, recent reports connect RpaA with the redox state of the pETC through interaction with ferredoxin (Fd) and thioredoxin (Trx) (Hanke et al., 2011; Kadowaki et al., 2015), and the activity of the latter act as a thiol redox switch to regulate the oligomeric state of the RpaA through redox-responsive cysteines (Ibrahim et al., 2022).

RpaA has also been implicated in central carbon metabolism and storage in cyanobacteria in ways that may indirectly contribute to the sensitivity of *rpaA* mutants to redox imbalances. Inefficient utilization or storage of primary products of photosynthesis is well documented to contribute to source/sink imbalances and photoinhibitory outcomes in a many green lineage species (Adams et al., 2014; Santos-Merino et al., 2021a). Conversely, remobilization of carbon from storage polymers (*i.e.*, glycogen) is essential to rapidly replenish photosynthetic carbohydrate intermediates to efficiently ‘reboot’ the Calvin-Benson-Bassham (CBB) cycle during environmental fluctuations, such as dark to light transitions (Makowka et al., 2020; Shinde et al., 2020). Previously, Δ*rpaA* mutants have been reported to inefficiently partition carbon towards glycogen under phototrophic conditions, despite the high levels of glycogen biosynthesis enzymes detected in this strain (Puszynska and O’Shea, 2017), and this may contribute to redox stress under light/dark cycles (Diamond et al., 2017). Here, we report that provided abundant organic carbon under mixotrophic growth, Δ*rpaA* mutants hyperaccumulate glycogen to levels above the typical physiological range (Figure 4). Our results demonstrate that Δ*rpaA* mutants are not deficient in glycogen synthesis capacity, and therefore may fail to provision adequate glycogen under phototrophic conditions due to other limitations (*e.g.*, misregulation of carbon partitioning and/or decreased CO_2_ fixation rates). Multiple lines of evidence therefore suggest that Δ*rpaA* mutants are unable to flexibly adapt the flux of carbon to match metabolic needs with changing environmental conditions. As a whole, Δ*rpaA* mutants may be especially overreactive to ROS due to the roles for RpaA in regulating metabolic redox pools, detoxifying ROS, and/or controlling alternative electron transport pathways that can remove excess reductant on the pETC.

Beyond their direct activity as potent oxidative compounds, ROS can also function as important signals to mediate a variety of cellular responses in photosynthetic organisms (Mullineaux et al., 2018). For example, in plants ROS have been proposed to be a primary signal of overreduction of the PQ pool (Mubarakshina and Ivanov, 2010), and H_2_O_2_ specifically has important signaling roles in a variety of plant processes including senescence, flowering, and biotic stress (Niu and Liao, 2016). In algae, H_2_O_2_-responsive pathways are specifically involved in the regulation of CCM, where it is proposed that ROS signals may serve as a proxy signal of imbalanced photosynthesis: activation of CCM processes may thereby help to rebalance the system and consequently suppresses further ROS formation (Choi et al., 2022). Of particular relevance to this study, it was recently reported that H_2_O_2_ (derived from photorespiratory processes or from direct chemical addition) can directly modulate the pyrenoid, inducing a more robust aggregation of rubisco into the pyrenoid core as well as a thicker and more defined pyrenoid starch sheath (Neofotis et al., 2021). Despite the lack of conservation in many of the structural components between carboxysomes and the algal pyrenoid, our data suggests that H_2_O_2_ derived from an overreduced pETC as a functionally conserved signal to coordinate downstream CCM (re)organization in cyanobacteria as well.

In natural environments, cyanobacteria experience changing environmental conditions (*e.g.*, high light, nutrient limitations) and must adaptively modulate core machinery to maintain robust photosynthesis (Stockenreiter et al., 2021). Cyanobacteria have evolved the ability to homeostatically regulate the redox status of its PQ pool, instead of functioning as a source of a regulatory signal (Schuurmans et al., 2014). Due to its central location between the two photosystems, the redox status of the PQ pool plays a pivotal role in sensing cellular status and in regulating photosynthetic capacity (Mullineaux and Allen, 1990). Particularly, the redox status of the PQ pool have regulatory roles related to photosystem composition (Fujita et al., 1987), state transitions (Mullineaux and Allen, 1990), and redistribution of respiratory complexes (Liu et al., 2012). Collectively, our data points to an additional regulatory role of the PQ pool as connected to the integrity and/or remodeling of the carbon fixation machinery under transitional periods where there may be a mismatch between the light and dark reactions. This control appears to involve RpaA, revealing a previously underappreciated function for the multi-layered regulation of both PQ and RpaA towards achieving energy balance in cyanobacteria.

## Acknowledgments

The Attune Cytpix, located in the MSU Flow Cytometry Core Facility, is supported by the Equipment Grants Program, award no. 2022-70410-38419, from the U.S. Department of Agriculture (USDA) National Institute of Food and Agriculture (NIFA). The authors thank Dr. Alicia Withrow and the Michigan State University Center for Advanced Microscopy for assistance with electron microscopy.

## 6. Funding

This work was primarily supported by the Department of Energy and Basic Energy Sciences Division (Grant: DE-FG02-91ER20021).

## Notes

### Competing Interest Statement

The authors have declared no competing interest.

## References

Abramson, B.W., Kachel, B., Kramer, D.M., and Ducat, D.C. (2016). Increased photochemical efficiency in cyanobacteria via an engineered sucrose sink. Plant & cell physiology 57, 2451–2460.

Adams, W.W., Muller, O., Cohu, C.M., and Demmig-Adams, B. (2014). Photosystem II efficiency and non-photochemical fluorescence quenching in the context of source-sink balance. In Non-Photochemical Quenching and Energy Dissipation in Plants, Algae and Cyanobacteria, B. Demmig-Adams, G. Garab, W. Adams Iii, and Govindjee, eds (Dordrecht: Springer Netherlands), pp. 503-529.

Alexova, R., Dang, T.C., Fujii, M., Raftery, M.J., Waite, T.D., Ferrari, B.C., and Neilan, B.A. (2016). Specific global responses to N and Fe nutrition in toxic and non-toxic *Microcystis aeruginosa*. Environ Microbiol 18, 401–413.

Allan, C., Burel, J.M., Moore, J., Blackburn, C., Linkert, M., Loynton, S., Macdonald, D., Moore, W.J., Neves, C., Patterson, A., Porter, M., Tarkowska, A., Loranger, B., Avondo, J., Lagerstedt, I., Lianas, L., Leo, S., Hands, K., Hay, R.T., Patwardhan, A., Best, C., Kleywegt, G.J., Zanetti, G., and Swedlow, J.R. (2012). OMERO: flexible, model-driven data management for experimental biology. Nat Methods 9, 245–253.

Ashby, M.K., and Mullineaux, C.W. (1999). Cyanobacterial *ycf27* gene products regulate energy transfer from phycobilisomes to photosystems I and II. FEMS Microbiol Lett 181, 253–260.

Baker, N.R., Harbinson, J., and Kramer, D.M. (2007). Determining the limitations and regulation of photosynthetic energy transduction in leaves. Plant Cell Environ 30, 1107–1125.

Barchewitz, T., Guljamow, A., Meissner, S., Timm, S., Henneberg, M., Baumann, O., Hagemann, M., and Dittmann, E. (2019). Non-canonical localization of RubisCO under high-light conditions in the toxic cyanobacterium *Microcystis aeruginosa* PCC7806. Environ Microbiol 21, 4836–4851.

Basalla, J.L., Ghalmi, M., Hoang, Y., Dow, R.E., and Vecchiarelli, A.G. (2024). An invariant C-terminal tryptophan in McdB mediates its interaction and positioning function with carboxysomes. Mol Biol Cell 35, ar107.

Beckmann, K., Messinger, J., Badger, M.R., Wydrzynski, T., and Hillier, W. (2009). On-line mass spectrometry: membrane inlet sampling. Photosynth Res 102, 511–522.

Bradley, L., Sipocz, B., Robitaille, T., Tollerud, E., Deil, C., Vinícius, Z., Barbary, K., Günther, H.M., Bostroem, A., Droettboom, M., Bray, E., Bratholm, L.A., Pickering, T.E., Craig, M., Pascual, S., Greco, J., Donath, A., Kerzendorf, W., Littlefair, S., Barentsen, G., D’Eugenio, F., and Weaver, B.A. (2016). Photutils: Photometry tools.

Burnap, R.L., Nambudiri, R., and Holland, S. (2013). Regulation of the carbon-concentrating mechanism in the cyanobacterium *Synechocystis* sp. PCC6803 in response to changing light intensity and inorganic carbon availability. Photosynth Res 118, 115-124.

Calzadilla, P.I., and Kirilovsky, D. (2020). Revisiting cyanobacterial state transitions. Photochem Photobiol Sci 19, 585–603.

Choi, B.Y., Kim, H., Shim, D., Jang, S., Yamaoka, Y., Shin, S., Yamano, T., Kajikawa, M., Jin, E., Fukuzawa, H., and Lee, Y. (2022). The *Chlamydomonas* bZIP transcription factor BLZ8 confers oxidative stress tolerance by inducing the carbon-concentrating mechanism. The Plant cell 34, 910–926.

Cohen, S.E., and Golden, S.S. (2015). Circadian rhythms in cyanobacteria. Microbiol Mol Biol Rev 79, 373–385.

Diamond, S., Jun, D., Rubin, B.E., and Golden, S.S. (2015). The circadian oscillator in *Synechococcus elongatus* controls metabolite partitioning during diurnal growth. Proceedings of the National Academy of Sciences of the United States of America 112, E1916–1925.

Diamond, S., Rubin, B.E., Shultzaberger, R.K., Chen, Y., Barber, C.D., and Golden, S.S. (2017). Redox crisis underlies conditional light-dark lethality in cyanobacterial mutants that lack the circadian regulator, RpaA. Proceedings of the National Academy of Sciences of the United States of America 114, E580–E589.

Dong, G., Yang, Q., Wang, Q., Kim, Y.I., Wood, T.L., Osteryoung, K.W., van Oudenaarden, A., and Golden, S.S. (2010). Elevated ATPase activity of KaiC applies a circadian checkpoint on cell division in *Synechococcus elongatus*. Cell 140, 529–539.

Ducat, D.C., Avelar-Rivas, J.A., Way, J.C., and Silver, P.A. (2012). Rerouting carbon flux to enhance photosynthetic productivity. Appl Environ Microbiol 78, 2660–2668.

Espinosa, J., Boyd, J.S., Cantos, R., Salinas, P., Golden, S.S., and Contreras, A. (2015). Cross-talk and regulatory interactions between the essential response regulator RpaB and cyanobacterial circadian clock output. Proceedings of the National Academy of Sciences of the United States of America 112, 2198–2203.

Fan, J., Zhou, D., Chen, C., Wu, J., and Wu, H. (2021). Reprogramming the metabolism of *Synechocystis* PCC 6803 by regulating the plastoquinone biosynthesis. Synth Syst Biotechnol 6, 351–359.

Foyer, C.H. (2018). Reactive oxygen species, oxidative signaling and the regulation of photosynthesis. Environ Exp Bot 154, 134–142.

Fujita, Y., Murakami, A., and Ohki, K. (1987). Regulation of photosystem composition in the cyanobacterial photosynthetic system: the regulation accurs in response to the redox state of the electron pool located between the two photosystems. Plant and Cell Physiology 28, 283–292.

Gründel, M., Scheunemann, R., Lockau, W., and Zilliges, Y. (2012). Impaired glycogen synthesis causes metabolic overflow reactions and affects stress responses in the cyanobacterium *Synechocystis* sp. PCC 6803. Microbiology 158, 3032-3043.

Hall, C.C., Cruz, J., Wood, M., Zegarac, R., DeMars, D., Carpenter, J., Kanazawa, A., and Kramer, D.M. (2013). Photosynthetic measurements with the Idea Spec: ani ntegrated diode emitter array spectrophotometer/fluorometer. In Photosynthesis Research for Food, Fuel and the Future, T. Kuang, C. Lu, and L. Zhang, eds (Berlin, Heidelberg: Springer Berlin Heidelberg), pp. 184-188.

Hanke, G.T., Satomi, Y., Shinmura, K., Takao, T., and Hase, T. (2011). A screen for potential ferredoxin electron transfer partners uncovers new, redox dependent interactions. Biochim Biophys Acta 1814, 366–374.

Hill, N.C., Tay, J.W., Altus, S., Bortz, D.M., and Cameron, J.C. (2020). Life cycle of a cyanobacterial carboxysome. Sci Adv 6, eaba1269.

Ibrahim, I.M., Rowden, S.J.L., Cramer, W.A., Howe, C.J., and Puthiyaveetil, S. (2022). Thiol redox switches regulate the oligomeric state of cyanobacterial Rre1, RpaA and RpaB response regulators. FEBS letters 596, 1533–1543.

Iijima, H., Shirai, T., Okamoto, M., Kondo, A., Hirai, M.Y., and Osanai, T. (2015). Changes in primary metabolism under light and dark conditions in response to overproduction of a response regulator RpaA in the unicellular cyanobacterium *Synechocystis* sp. PCC 6803. Front Microbiol 6, 888.

Ivleva, N.B., Gao, T., LiWang, A.C., and Golden, S.S. (2006). Quinone sensing by the circadian input kinase of the cyanobacterial circadian clock. Proceedings of the National Academy of Sciences of the United States of America 103, 17468–17473.

Kadowaki, T., Nishiyama, Y., Hisabori, T., and Hihara, Y. (2015). Identification of OmpR-family response regulators interacting with thioredoxin in the Cyanobacterium *Synechocystis* sp. PCC 6803. PloS one 10, e0119107.

Khorobrykh, S., Tsurumaki, T., Tanaka, K., Tyystjarvi, T., and Tyystjarvi, E. (2020). Measurement of the redox state of the plastoquinone pool in cyanobacteria. FEBS letters 594, 367–375.

Kim, P., Kaur, M., Jang, H.I., and Kim, Y.I. (2020). The circadian clock-a molecular tool for survival in cyanobacteria. Life 10.

Kim, Y.I., Vinyard, D.J., Ananyev, G.M., Dismukes, G.C., and Golden, S.S. (2012). Oxidized quinones signal onset of darkness directly to the cyanobacterial circadian oscillator. Proceedings of the National Academy of Sciences of the United States of America 109, 17765–17769.

Kirilovsky, D., and Kerfeld, C.A. (2016). Cyanobacterial photoprotection by the orange carotenoid protein. Nat Plants 2, 16180.

Krieger-Liszkay, A., and Shimakawa, G. (2022). Regulation of the generation of reactive oxygen species during photosynthetic electron transport. Biochem Soc Trans 50, 1025–1034.

Labiosa, R.G., Arrigo, K.R., Tu, C.J., Bhaya, D., Bay, S., Grossman, A.R., and Shrager, J. (2006). Examination of diel changes in global transcript accumulation in *Synechocystis* (cyanobacteria). Journal of Phycology 42, 622–636.

Latifi, A., Ruiz, M., and Zhang, C.C. (2009). Oxidative stress in cyanobacteria. FEMS Microbiol Rev 33, 258–278.

Lea-Smith, D.J., Ross, N., Zori, M., Bendall, D.S., Dennis, J.S., Scott, S.A., Smith, A.G., and Howe, C.J. (2013). Thylakoid terminal oxidases are essential for the cyanobacterium *Synechocystis* sp. PCC 6803 to survive rapidly changing light intensities. Plant physiology 162, 484-495.

Liu, L.N., Bryan, S.J., Huang, F., Yu, J., Nixon, P.J., Rich, P.R., and Mullineaux, C.W. (2012). Control of electron transport routes through redox-regulated redistribution of respiratory complexes. Proceedings of the National Academy of Sciences of the United States of America 109, 11431–11436.

Long, B.M., Badger, M.R., Whitney, S.M., and Price, G.D. (2007). Analysis of carboxysomes from *Synechococcus* PCC7942 reveals multiple Rubisco complexes with carboxysomal proteins CcmM and CcaA. J Biol Chem 282, 29323–29335.

Lucius, S., and Hagemann, M. (2024). The primary carbon metabolism in cyanobacteria and its regulation. Front Plant Sci 15, 1417680.

MacCready, J.S., Hakim, P., Young, E.J., Hu, L., Liu, J., Osteryoung, K.W., Vecchiarelli, A.G., and Ducat, D.C. (2018). Protein gradients on the nucleoid position the carbon-fixing organelles of cyanobacteria. eLife 7.

Makowka, A., Nichelmann, L., Schulze, D., Spengler, K., Wittmann, C., Forchhammer, K., and Gutekunst, K. (2020). Glycolytic shunts replenish the Calvin-Benson-Bassham cycle as anaplerotic reactions in cyanobacteria. Mol Plant 13, 471–482.

Markson, J.S., Piechura, J.R., Puszynska, A.M., and O’Shea, E.K. (2013). Circadian control of global gene expression by the cyanobacterial master regulator RpaA. Cell 155, 1396–1408.

Martins, B.M.C., Tooke, A.K., Thomas, P., and Locke, J.C.W. (2018). Cell size control driven by the circadian clock and environment in cyanobacteria. Proceedings of the National Academy of Sciences of the United States of America 115, E11415–E11424.

Mironov, K.S., Sinetova, M.A., Shumskaya, M., and Los, D.A. (2019). Universal molecular triggers of stress responses in cyanobacterium *Synechocystis*. Life 9.

Mubarakshina, M.M., and Ivanov, B.N. (2010). The production and scavenging of reactive oxygen species in the plastoquinone pool of chloroplast thylakoid membranes. Physiol Plant 140, 103–110.

Mullineaux, C.W., and Allen, J.F. (1990). State 1-State 2 transitions in the cyanobacterium *Synechococcus* 6301 are controlled by the redox state of electron carriers between Photosystems I and II. Photosynth Res 23, 297–311.

Mullineaux, P.M., Exposito-Rodriguez, M., Laissue, P.P., and Smirnoff, N. (2018). ROS-dependent signalling pathways in plants and algae exposed to high light: Comparisons with other eukaryotes. Free radical biology & medicine 122, 52–64.

Muth-Pawlak, D., Kreula, S., Gollan, P.J., Huokko, T., Allahverdiyeva, Y., and Aro, E.M. (2022). Patterning of the autotrophic, mixotrophic, and heterotrophic proteomes of oxygen-evolving cyanobacterium *Synechocystis* sp. PCC 6803. Front Microbiol 13, 891895.

Nakajima, M., Imai, K., Ito, H., Nishiwaki, T., Murayama, Y., Iwasaki, H., Oyama, T., and Kondo, T. (2005). Reconstitution of circadian oscillation of cyanobacterial KaiC phosphorylation *in vitro*. Science 308, 414–415.

Neofotis, P., Temple, J., Tessmer, O.L., Bibik, J., Norris, N., Pollner, E., Lucker, B., Weraduwage, S.M., Withrow, A., Sears, B., Mogos, G., Frame, M., Hall, D., Weissman, J., and Kramer, D.M. (2021). The induction of pyrenoid synthesis by hyperoxia and its implications for the natural diversity of photosynthetic responses in *Chlamydomonas*. eLife 10.

Nikkanen, L., Solymosi, D., Jokel, M., and Allahverdiyeva, Y. (2021). Regulatory electron transport pathways of photosynthesis in cyanobacteria and microalgae: Recent advances and biotechnological prospects. Physiol Plant 173, 514–525.

Niu, L., and Liao, W. (2016). Hydrogen peroxide signaling in plant development and abiotic responses: crosstalk with nitric oxide and calcium. Front Plant Sci 7, 230.

Pattanayak, G.K., Phong, C., and Rust, M.J. (2014). Rhythms in energy storage control the ability of the cyanobacterial circadian clock to reset. Current biology : CB 24, 1934–1938.

Perelman, A., Uzan, A., Hacohen, D., and Schwarz, R. (2003). Oxidative stress in *Synechococcus* sp. strain PCC 7942: various mechanisms for H_2_O_2_ detoxification with different physiological roles. J Bacteriol 185, 3654–3660.

Porra, R.J., Thompson, W.A., and Kriedemann, P.E. (1989). Determination of accurate extinction coefficients and simultaneous equations for assaying chlorophylls *a* and *b* extracted with four different solvents: verification of the concentration of chlorophyll standards by atomic absorption spectroscopy. Biochimica et Biophysica Acta (BBA) - Bioenergetics 975, 384–394.

Pospisil, P. (2016). Production of reactive oxygen species by Photosystem II as a response to light and temperature stress. Front Plant Sci 7, 1950.

Puszynska, A.M., and O’Shea, E.K. (2017). Switching of metabolic programs in response to light availability is an essential function of the cyanobacterial circadian output pathway. eLife 6.

Qiao, C., Zhang, M., Luo, Q., and Lu, X. (2019). Identification of two two-component signal transduction mutants with enhanced sucrose biosynthesis in *Synechococcus elongatus* PCC 7942. J Basic Microbiol 59, 465–476.

Rillema, R., Hoang, Y., MacCready, J.S., and Vecchiarelli, A.G. (2021). Carboxysome mispositioning alters growth, morphology, and Rubisco level of the cyanobacterium *Synechococcus elongatus* PCC 7942. mBio 12, e0269620.

Rohnke, B.A., Singh, S.P., Pattanaik, B., and Montgomery, B.L. (2018). RcaE-dependent regulation of carboxysome structural proteins has a central role in environmental determination of carboxysome morphology and Aabundance in *Fremyella diplosiphon*. mSphere 3.

Sacksteder, C.A., and Kramer, D.M. (2000). Dark-interval relaxation kinetics (DIRK) of absorbance changes as a quantitative probe of steady-state electron transfer. Photosynth Res 66, 145–158.

Sakkos, J.K., Hernandez-Ortiz, S., Osteryoung, K.W., and Ducat, D.C. (2021). Orthogonal degron system for controlled protein degradation in cyanobacteria. ACS Synth Biol 10, 1667–1681.

Santos-Merino, M., Singh, A.K., and Ducat, D.C. (2021a). Sink engineering in photosynthetic microbes. In Cyanobacteria Biotechnology, pp. 171-209.

Santos-Merino, M., Sakkos, J.K., Singh, A.K., and Ducat, D.C. (2024). Coordination of carbon partitioning and photosynthesis by a two-component signaling network in *Synechococcus elongatus* PCC 7942. Metabolic engineering 81, 38–52.

Santos-Merino, M., Torrado, A., Davis, G.A., Rottig, A., Bibby, T.S., Kramer, D.M., and Ducat, D.C. (2021b). Improved photosynthetic capacity and photosystem I oxidation via heterologous metabolism engineering in cyanobacteria. Proceedings of the National Academy of Sciences of the United States of America 118.

Schuurmans, J.M., Brinkmann, B.W., Makower, A.K., Dittmann, E., Huisman, J., and Matthijs, H.C.P. (2018). Microcystin interferes with defense against high oxidative stress in harmful cyanobacteria. Harmful Algae 78, 47–55.

Schuurmans, R.M., Schuurmans, J.M., Bekker, M., Kromkamp, J.C., Matthijs, H.C., and Hellingwerf, K.J. (2014). The redox potential of the plastoquinone pool of the cyanobacterium *Synechocystis* species strain PCC 6803 is under strict homeostatic control. Plant physiology 165, 463–475.

Shinde, S., Zhang, X., Singapuri, S.P., Kalra, I., Liu, X., Morgan-Kiss, R.M., and Wang, X. (2020). Glycogen metabolism supports photosynthesis start through the oxidative pentose phosphate pathway in cyanobacteria. Plant physiology 182, 507–517.

Singh, A.K., Santos-Merino, M., Sakkos, J.K., Walker, B.J., and Ducat, D.C. (2022). Rubisco regulation in response to altered carbon status in the cyanobacterium *Synechococcus elongatus* PCC 7942. Plant physiology 189, 874–888.

Stockenreiter, M., Isanta Navarro, J., Buchberger, F., and Stibor, H. (2021). Community shifts from eukaryote to cyanobacteria dominated phytoplankton: The role of mixing depth and light quality. Freshwater Biology 66, 2145–2157.

Stringer, C., Wang, T., Michaelos, M., and Pachitariu, M. (2021). Cellpose: a generalist algorithm for cellular segmentation. Nat Methods 18, 100–106.

Sun, Y., Huang, F., Dykes, G.F., and Liu, L.N. (2020). Diurnal regulation of in vivo localization and CO_2_-fixing activity of carboxysomes in *Synechococcus elongatus* PCC 7942. Life 10.

Sun, Y., Wollman, A.J.M., Huang, F., Leake, M.C., and Liu, L.N. (2019). Single-organelle quantification reveals stoichiometric and structural variability of carboxysomes dependent on the environment. The Plant cell 31, 1648–1664.

Sun, Y., Casella, S., Fang, Y., Huang, F., Faulkner, M., Barrett, S., and Liu, L.N. (2016). Light modulates the biosynthesis and organization of cyanobacterial carbon fixation machinery through photosynthetic electron flow. Plant physiology 171, 530–541.

Sunil, B., Talla, S.K., Aswani, V., and Raghavendra, A.S. (2013). Optimization of photosynthesis by multiple metabolic pathways involving interorganelle interactions: resource sharing and ROS maintenance as the bases. Photosynth Res 117, 61–71.

Takai, N., Nakajima, M., Oyama, T., Kito, R., Sugita, C., Sugita, M., Kondo, T., and Iwasaki, H. (2006). A KaiC-associating SasA-RpaA two-component regulatory system as a major circadian timing mediator in cyanobacteria. Proceedings of the National Academy of Sciences of the United States of America 103, 12109–12114.

Tan, L.R., Cao, Y.Q., Li, J.W., Xia, P.F., and Wang, S.G. (2022). Transcriptomics and metabolomics of engineered *Synechococcus elongatus* during photomixotrophic growth. Microbial cell factories 21, 31.

Taton, A., Erikson, C., Yang, Y., Rubin, B.E., Rifkin, S.A., Golden, J.W., and Golden, S.S. (2020). The circadian clock and darkness control natural competence in cyanobacteria. Nature communications 11, 1688.

Wang, B., Zuniga, C., Guarnieri, M.T., Zengler, K., Betenbaugh, M., and Young, J.D. (2023). Metabolic engineering of *Synechococcus elongatus* 7942 for enhanced sucrose biosynthesis. Metabolic engineering 80, 12–24.

Yang, Q., Pando, B.F., Dong, G., Golden, S.S., and van Oudenaarden, A. (2010). Circadian gating of the cell cycle revealed in single cyanobacterial cells. Science 327, 1522–1526.

