## Supplementary material for "Plastoquinone redox status influences carboxysome integrity via a RpaA- and ROS-dependent regulatory network"

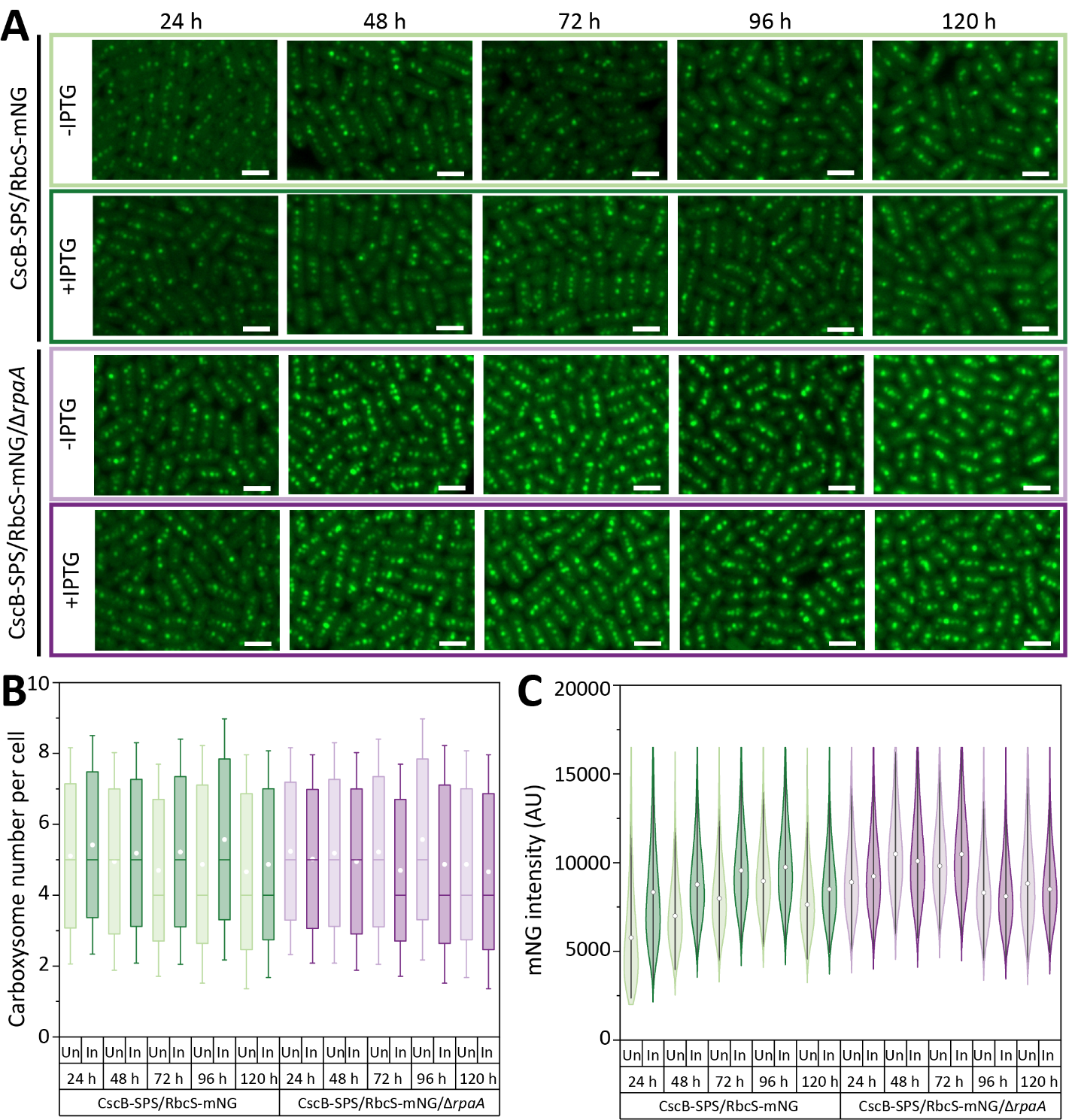

**Supplemental Figure S1. Δ*rpaA* mutants do not reorganize carboxysomes in response to sucrose export. (A)** Time-course of carboxysomes visualized via the RbcS-mNG reporter in the strain CscB-SPS^export^ in the presence/absence of RpaA. Scale bar: 2 µm. **(B)** Carboxysome number per cell in the strain CscB-SPS^export^ in the presence/absence of RpaA. White squares represent the media, the horizontal lines indicate the median and the whisker bars represent the standard deviation. **(C)** Carboxysome puncta mNG fluorescence intensity in the strain CscB-SPS^export^ in the presence/absence of RpaA. White circles represent the median and the bars represent the 95% confidence interval. **(B, C)** Un, uninduced (-IPTG); In, induced (+IPTG).

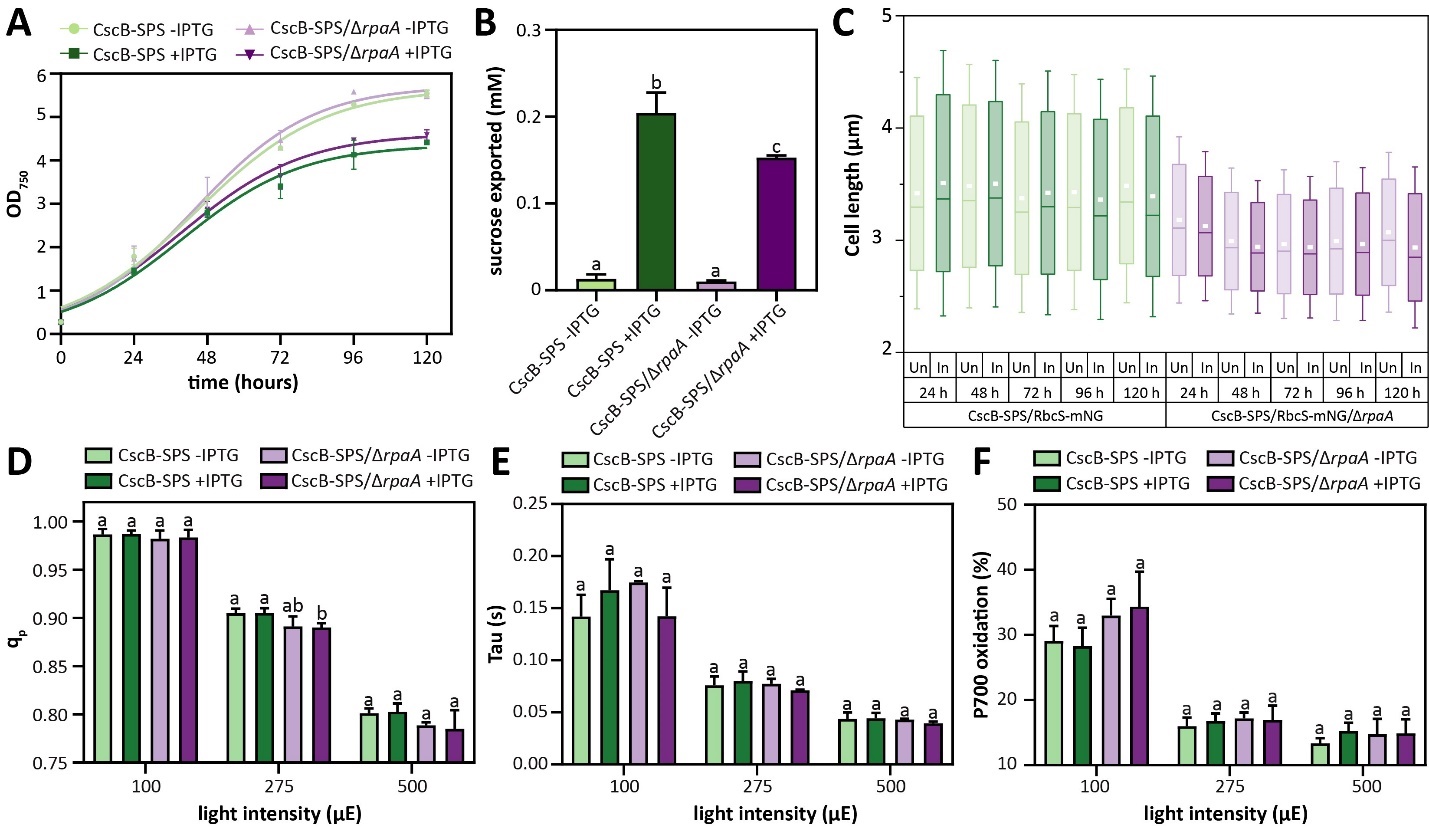

**Supplementary Figure S2. Knockout of RpaA does not globally disrupt cell physiology under constant light (A)** Growth curves following induction of sucrose export in the strain CscB-SPS^export^ in the presence/absence of RpaA. Averages of ≥3 independent biological replicates are shown ± SD. **(B)** Sucrose exported at 24 hours following induction of sucrose production and export pathways in the strain CscB-SPS^export^ in the presence/absence of RpaA. **(C)** Cell size under sucrose export conditions of the strain CscB-SPS^export^ in the presence/absence of RpaA. White squares represent the median, the horizontal lines indicate the median and the whisker bars represent the standard deviation. Un, uninduced (-IPTG); In, induced (+IPTG). **(D)** q_p_ values measured at three different light intensities 24 hours after the induction of sucrose export in the strain CscB-SPS^export^ in the presence/absence of RpaA. **(E)** Tau values, expressed in seconds, measured at three different light intensities 24 hours after the induction of sucrose export in the strain CscB-SPS^export^ in the presence/absence of RpaA. **(F)** P700 oxidation levels, expressed as a percentage, measured at three different light intensities 24 h after the induction of sucrose export in the strain CscB-SPS^export^ in the presence/absence of RpaA. **(B, D, E, F)** Averages of ≥3 independent biological replicates are shown + SD. Significance was calculated by one-way ANOVA followed by Tukey’s multiple comparison test. Data points labeled with different letters are significantly different (*P* < 0.05).

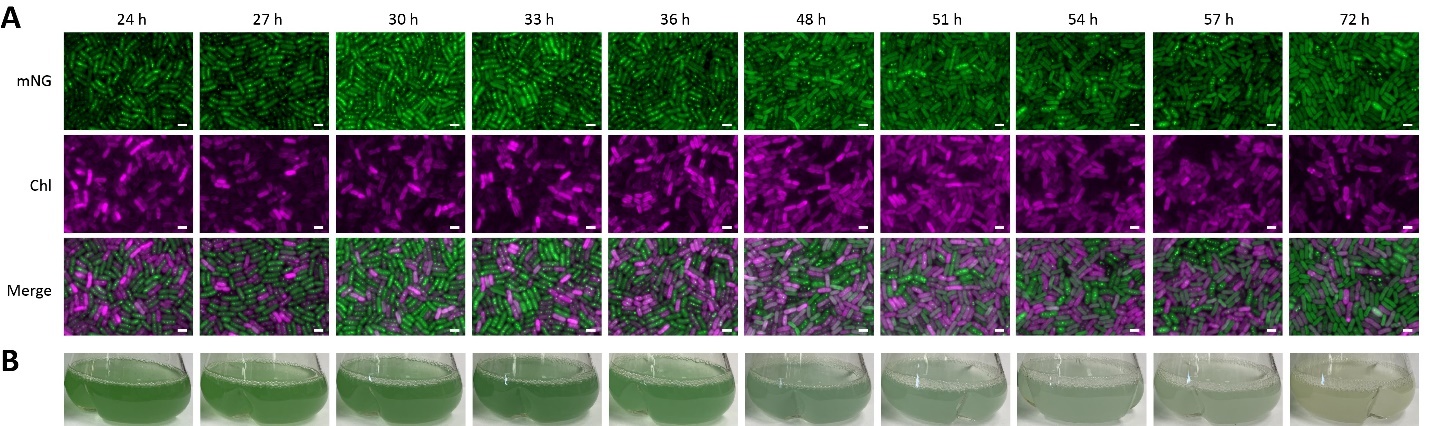
**Supplementary Figure S3. Time-course of carboxysome disassembly in the CscB/RbcS-mNG/Δ*rpaA* strain under mixotrophic conditions. (A)** Time-course of fluorescence microscope images showing the disassembly of carboxysomes. **(B)** Time-course of flask images to show culture appearance and loss of pigmentation.

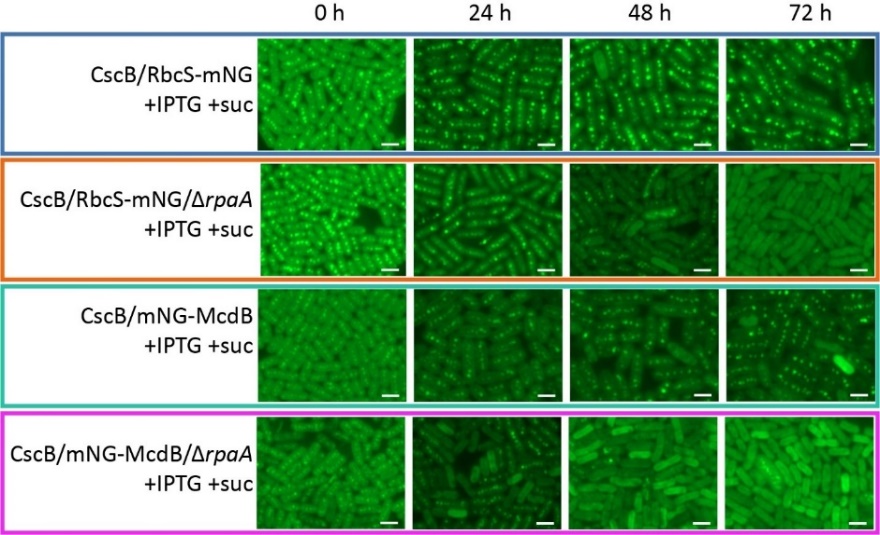

**Supplementary Figure S4. The outer carboxysome component McdB is delocalized prior to the loss of core rubisco puncta following onset of mixotrophic conditions in Δ*rpaA* cells.** Time-course of carboxysome status under mixotrophic conditions in the strain CscB^import^ in the presence/absence of RpaA by tracking either RbcS-mNG or mNG-McdB. Scale bar: 2 µm.

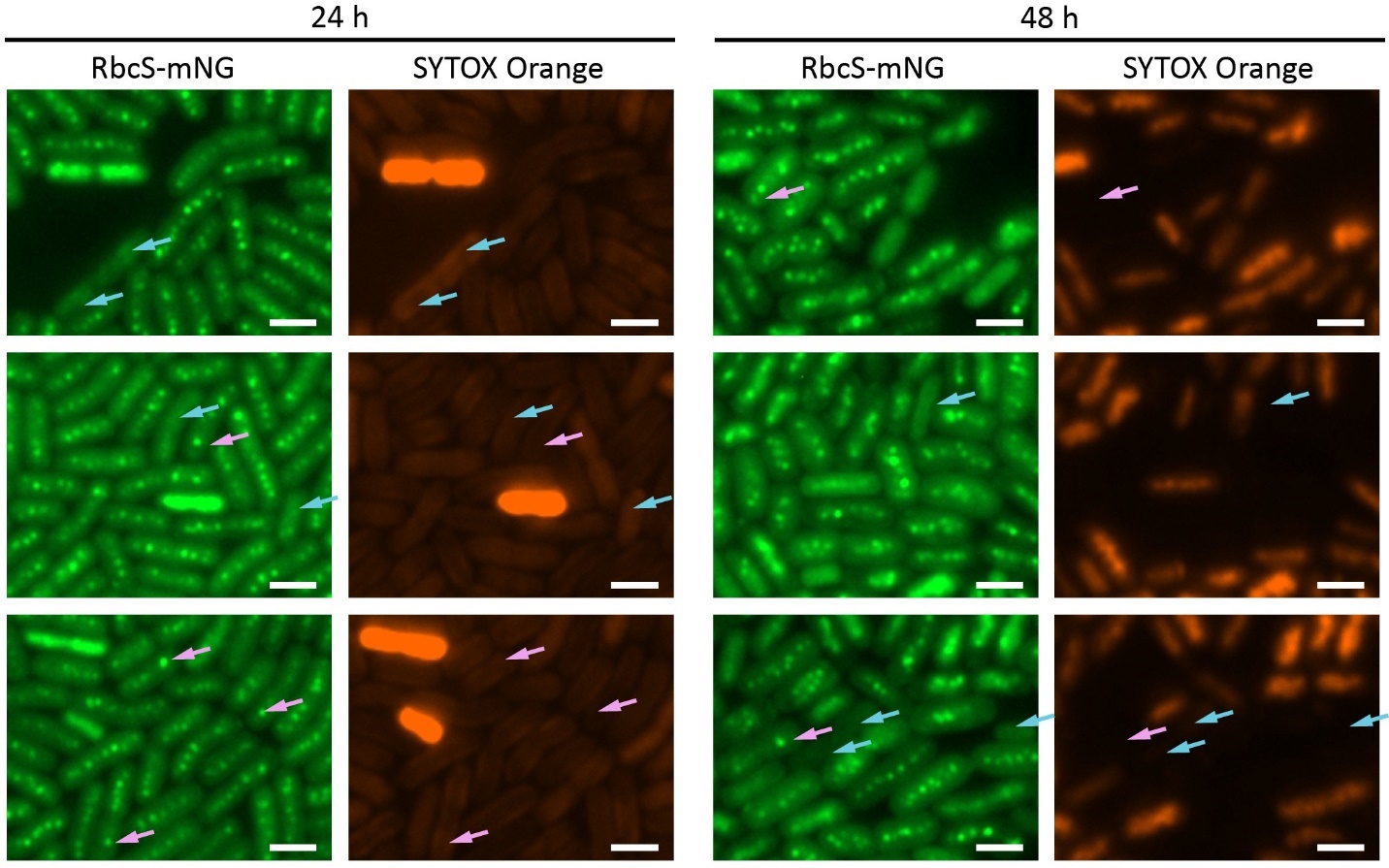

**Supplementary Figure S5**. **Mobilization of carboxysome to the cell poles and carboxysome breakdown happens prior cell death.** Detailed fluorescence microscopy images at 24 and 48 hours of cells stained with SYTOX-Orange and carboxysomes labeled with mNG in response to sucrose feeding in the strain CscB^import^ in the absence of RpaA. Scale bar: 2 µm. Blue arrows indicate cells with no carboxysomes and SYTOX-Orange negative (alive cells). Pink arrows indicate cells which carboxysomes have been mobilized to the cell poles to be potentially disassembled.

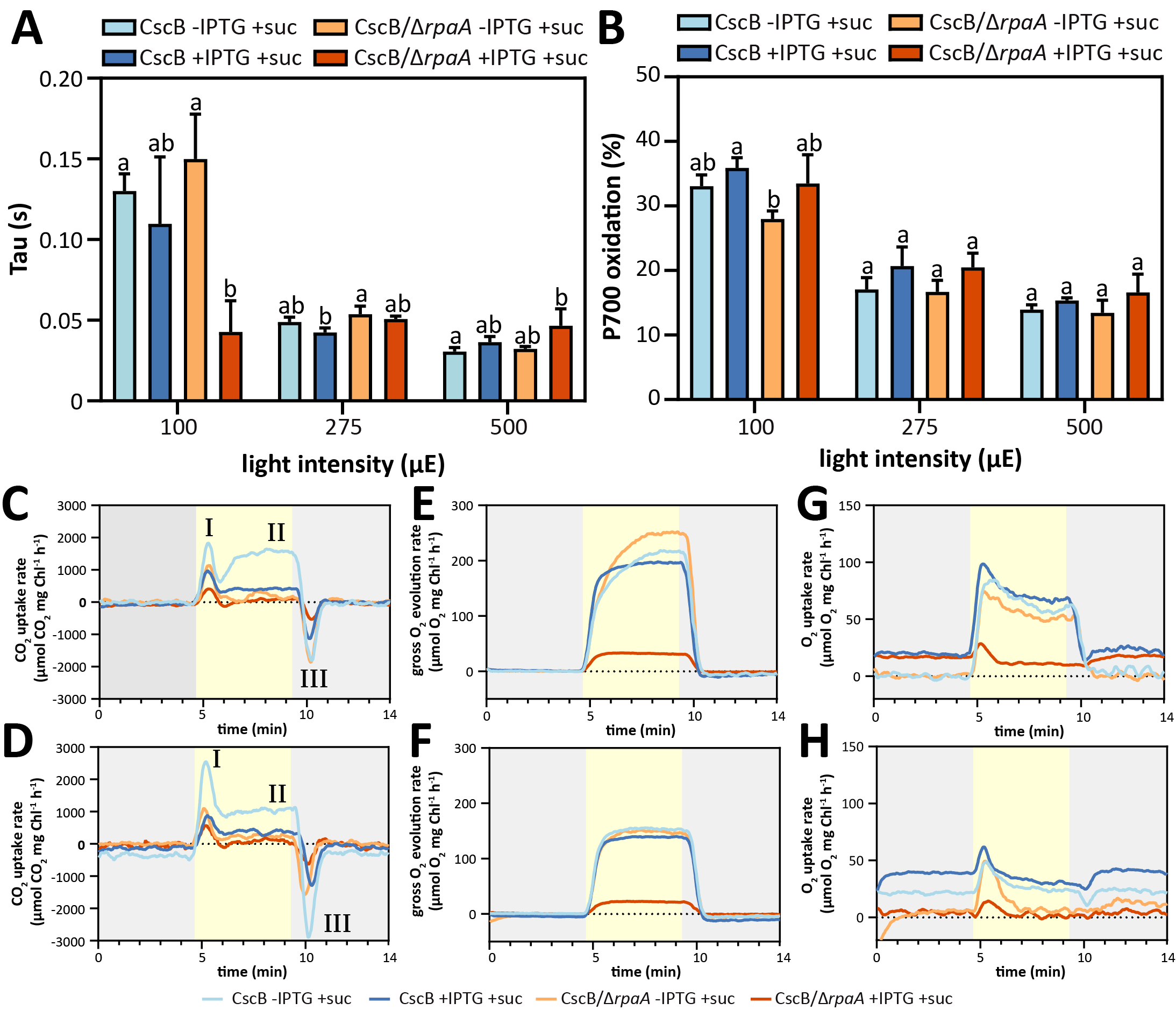

**Supplementary Figure S6. Changes in photosynthesis and O_2_ and CO_2_ fluxes associated with mixotrophic conditions in Δ*rpaA*. (A)** Tau values. **(B)** P700 oxidation levels, expressed as a percentage. CO_2_ exchange rates in the strains CscB/RbcS-mNG and CscB/RbcS-mNG/Δ*rpaA* under photoautotrophic and mixotrophic conditions **(C)** at 24 hours and **(D)** at 48 hours post-induction. Gross O_2_ evolution rates under photoautotrophic and mixotrophic conditions **(E)** at 24 hours and **(F)** at 48 hours post-induction. O_2_ reduction rates under photoautotrophic and mixotrophic conditions **(E)** at 24 hours and **(F)** at 48 hours post-induction. **(A, C)** Averages of ≥3 independent biological replicates are shown + SD. Significance was calculated by one-way ANOVA followed by Tukey’s multiple comparison test. Data points labeled with different letters are significantly different (*P* < 0.05). **(C, D, E, F, G, H)** Dark periods are indicated by grey rectangles. Yellow rectangles represent exposure to a light intensity of 500 μmol photons m^−2^ s^−1^. **(C, D)** Phase I **-** upon the onset of light, the carbon concentration mechanism (CCM) triggered an initial concentration step absorbing high amounts of CO_2_; Phase II - during the rest of the light period, consumption reaches a steady state; Phase III - when the lights are switched off, CCM activity ceases, leading to the release and efflux of the intracellular inorganic carbon pool back into the media. Dashed lines indicate the compensation point of CO_2_ fixation (uptake rate equals respiratory rate). **(E, F, G, H)** Dashed lines indicate the light compensation point (oxygen produced by photosynthesis equals oxygen consumed by respiration).

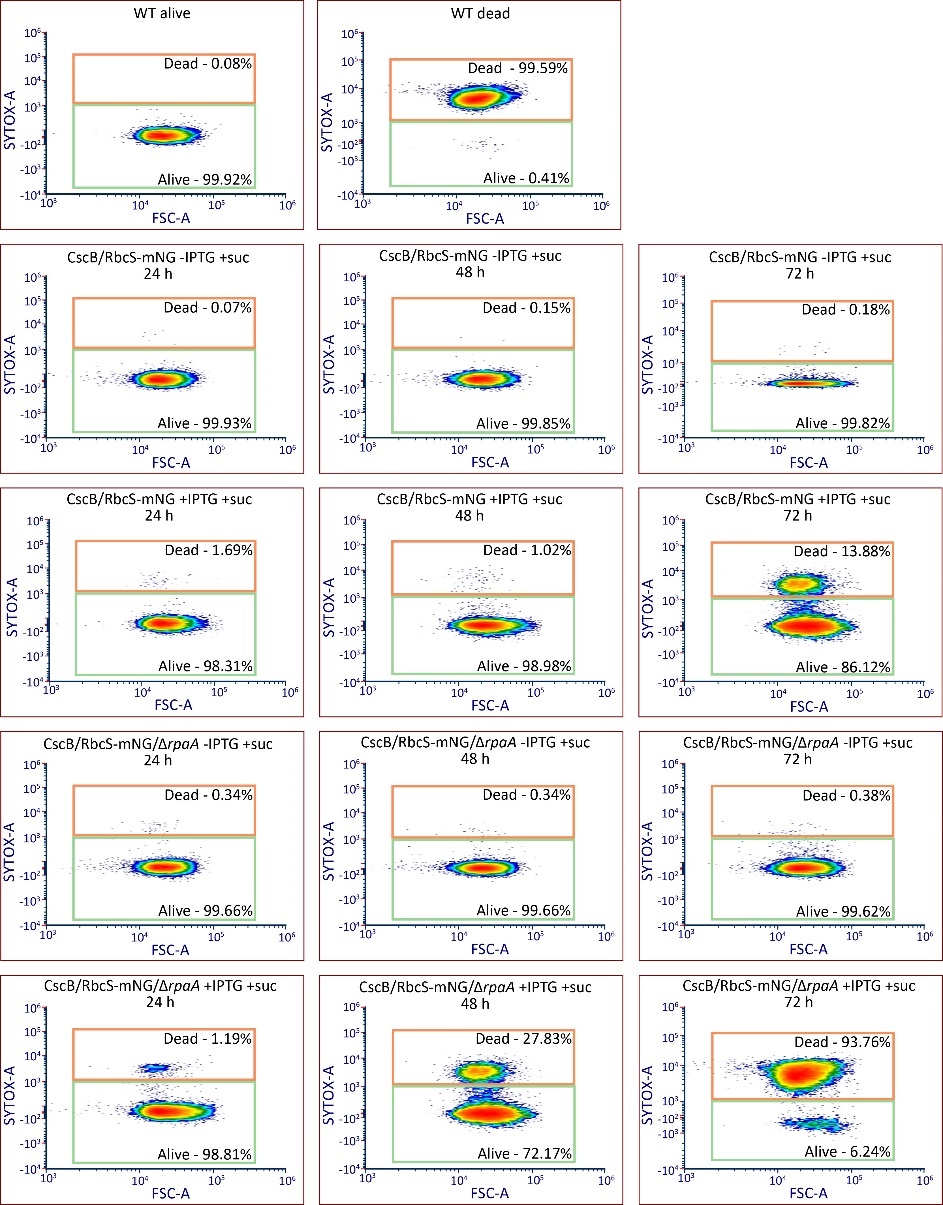

**Supplementary Figure S7. Dot plots of SYTOX blue flow cytometry revealed a negative effect of sucrose feeding in the Δ*rpaA* mutant.** Alive, SYTOX-negative cells; Dead, SYTOX-positive cells; FSC-A, forward scatter area; SYTOX-A, SYTOX blue area.

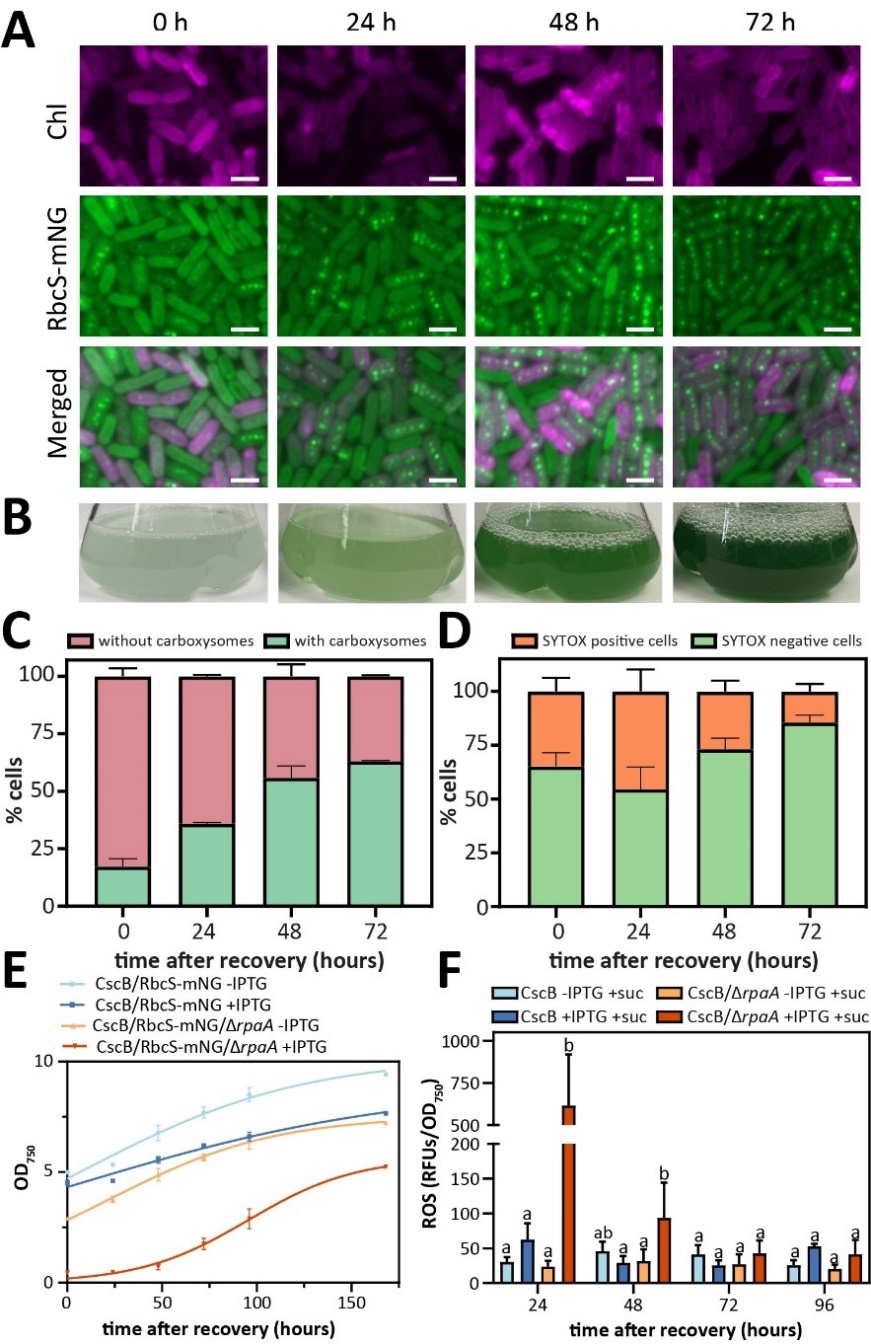

**Supplementary Figure S8. Sucrose removal from media allows Δ*rpaA* cells to recover carboxysomes, viability and growth and reduces ROS. (A)** Time-course of carboxysome recovery after removal of the sucrose from the media in the strain CscB^import^ in the absence of RpaA. Scale bar: 2 µm. **(B)** Time-course of flask images to show culture appearance and recovery of pigmentation. **(C)** Quantification of the number of carboxysomes over time after sucrose removal from the media. **(D)** Percentage of SYTOX-positive cells after removal of the sucrose from the media in the strain CscB^import^ in the absence of RpaA. **(E)** Growth curves of the strain CscB^import^ in the presence/absence of RpaA in response to sucrose removal from the media. Averages of ≥3 independent biological replicates are shown ± SD. **(F)** Quantification of cellular ROS accumulation measured by H_2_DCFDA fluorescence at different time points following removal of sucrose from the media of the CscB^import^ strain in the absence of RpaA. Averages of ≥3 independent biological replicates are shown + SD. Significance was calculated by one-way ANOVA followed by Tukey’s multiple comparison test. Data points labeled with different letters are significantly different (*P* < 0.05). **(C, D)** Averages of ≥3 independent biological replicates are shown ± SD.

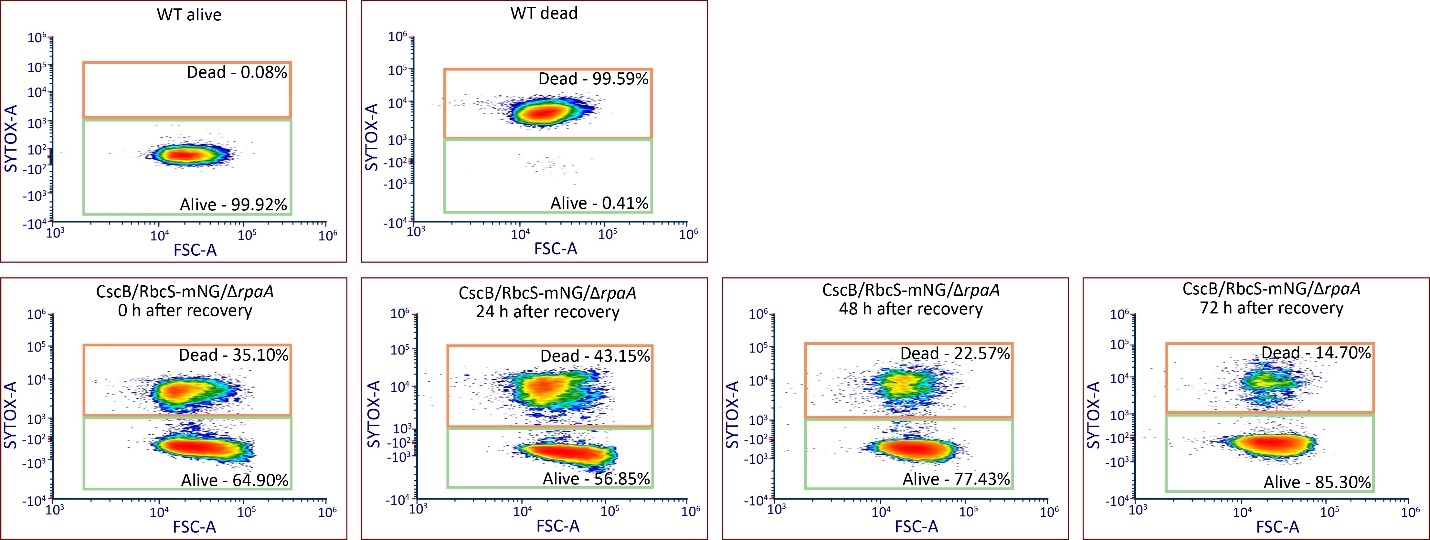
**Supplementary Figure S9. Dot plots of SYTOX blue flow cytometry revealed the recovery of the viability in the Δ*rpaA* mutant after sucrose removal from the media.** Alive, SYTOX-negative cells; Dead, SYTOX-positive cells; FSC-A, forward scatter area; SYTOX-A, SYTOX blue area.

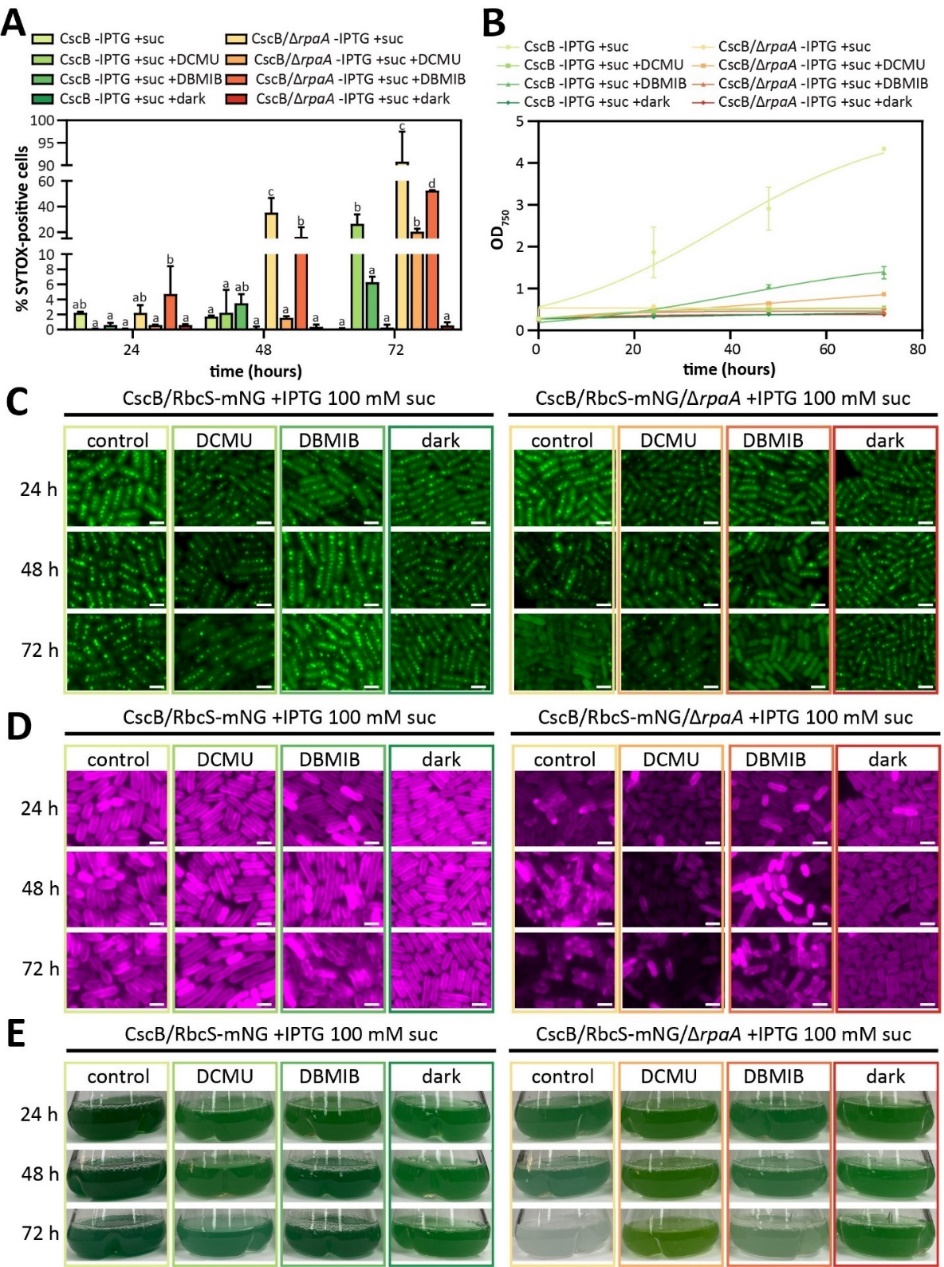

**Supplementary Figure S10. Effect of photosynthesis inhibitors and darkness treatment in the population viability, growth, carboxysomes, chlorophyll and culture aspect. (A)** Percentage of SYTOX-positive cells (dead cells) under mixotrophic conditions in the CscB^import^ strain in the presence/absence of RpaA following the addition of photosynthesis inhibitors or growing the cells in darkness. Averages of ≥3 independent biological replicates are shown + SD. Significance was calculated by one-way ANOVA followed by Tukey’s multiple comparison test. Data points labeled with different letters are significantly different (*P* < 0.05). **(B)** Growth curves under mixotrophic conditions of the CscB^import^ strain in the presence/absence of RpaA following the addition of photosynthesis inhibitors or growing the cells in darkness. Averages of ≥3 independent biological replicates are shown ± SD. **(C)** Carboxysome status under mixotrophic conditions of the CscB^import^ strain in the presence/absence of RpaA following the addition of photosynthesis inhibitors or growing the cells in darkness by tracking RbcS-mNG. Scale bar: 2 µm. **(D)** Chlorophyll changes under mixotrophic conditions of the CscB^import^ strain in the presence/absence of RpaA following the addition of photosynthesis inhibitors or growing the cells in darkness by tracking the red channel. Scale bar: 2 µm. **(E)** Changes in the appearance of the cultures after treatment with photosynthesis inhibitors or growing the cells in darkness under mixotrophic conditions of the CscB^import^ strain in the presence/absence of RpaA.

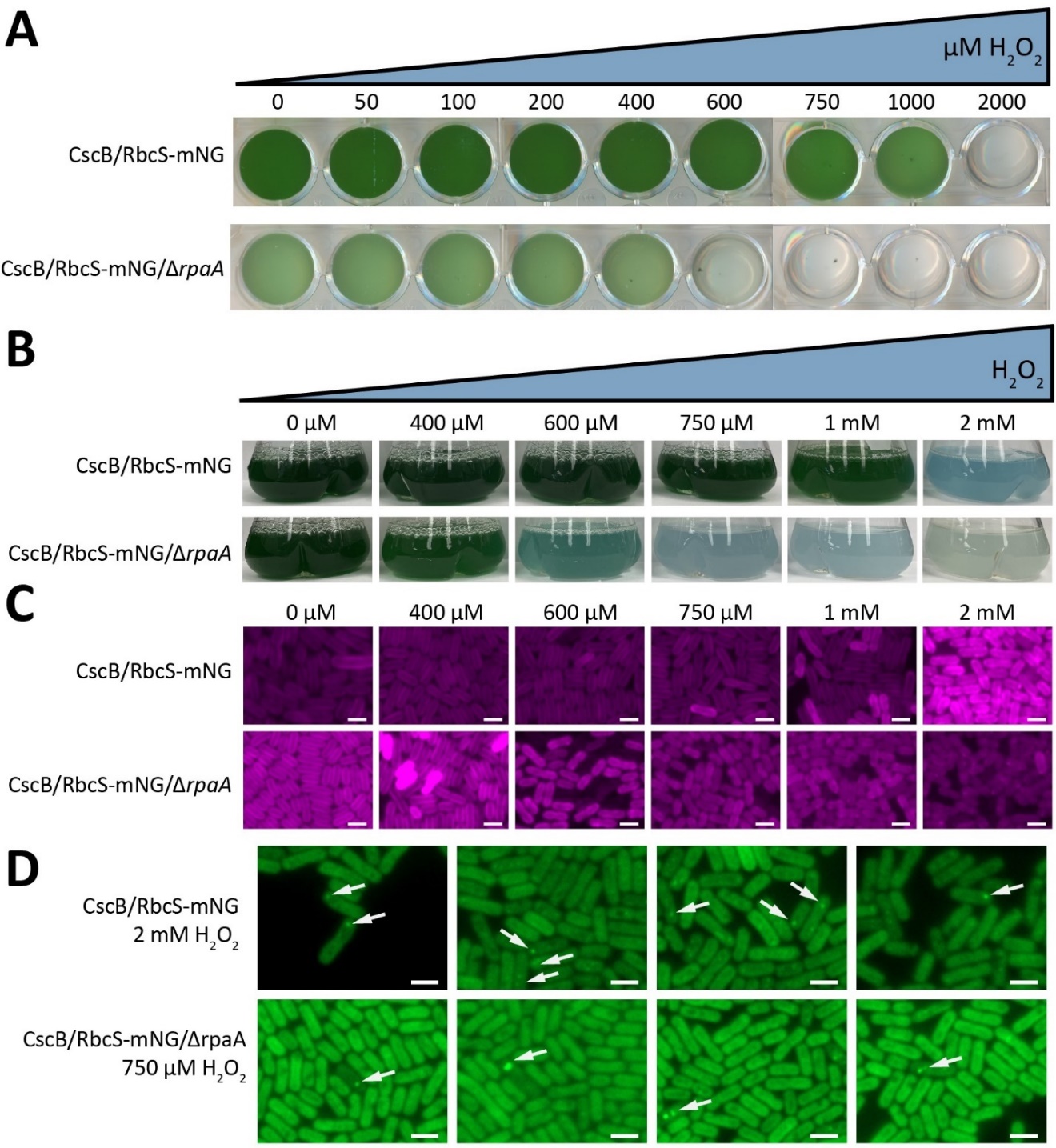
**Supplementary Figure S11. Effects of H_2_O_2_ treatment. (A)** Multiwell plate assay to estimate the tolerance to H_2_O_2_ of the strains CscB/RbcS-mNG and CscB/RbcS-mNG/Δ*rpaA* without sucrose feeding activated after 24 h of H_2_O_2_ exposure. **(B)** Changes in the appearance of the cultures after treatment with H_2_O_2_. CscB/RbcS-mNG and CscB/RbcS-mNG/Δ*rpaA* strains were grown in the presence of different concentrations of H_2_O_2_ for 24 h and changes in the color of the cultures were observed turning from green to blue. **(C)** Chlorophyll changes in response to different concentrations of H_2_O_2_ in the strain CscB^import^ in the presence/absence of RpaA after 24 h of H_2_O_2_ exposure by tracking red channel in the fluorescence microscope. Scale bar: 2 µm. **(D)** Detailed images showing the polar localization of carboxysomes in response to H_2_O_2_ treatment (white arrow). Scale bar: 2 µm.

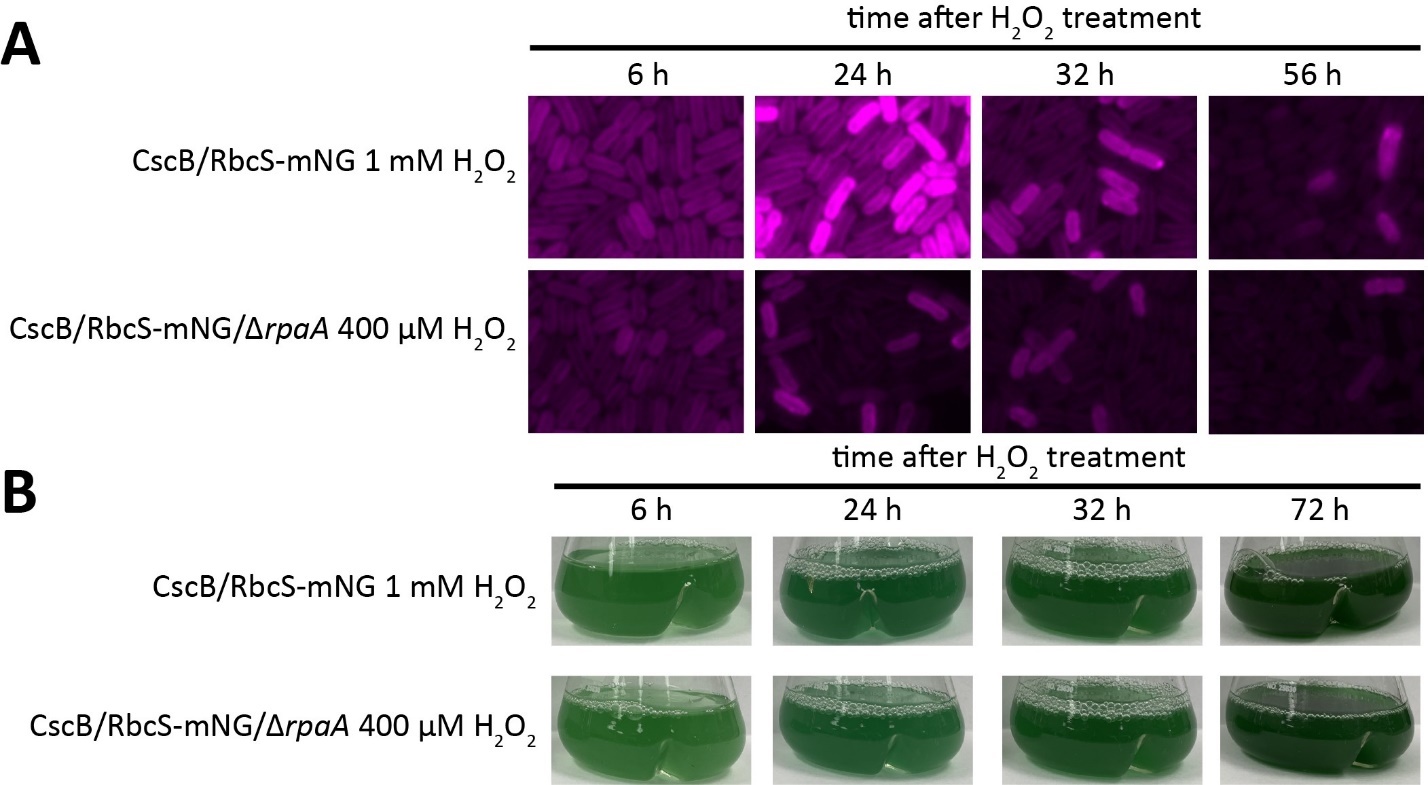

**Supplementary Figure S12. Effect of sublethal concentrations of H_2_O_2_ in the chlorophyll and culture aspect. (A)** Chlorophyll changes at different points after treatment with sublethal concentrations of H_2_O_2_ in the CscB^import^ strain in the presence/absence of RpaA by tracking the red channel. Scale bar: 2 µm. **(E)** Changes in the appearance of the cultures after treatment with sublethal concentrations of H_2_O_2_ in the CscB^import^ strain in the presence/absence of RpaA.

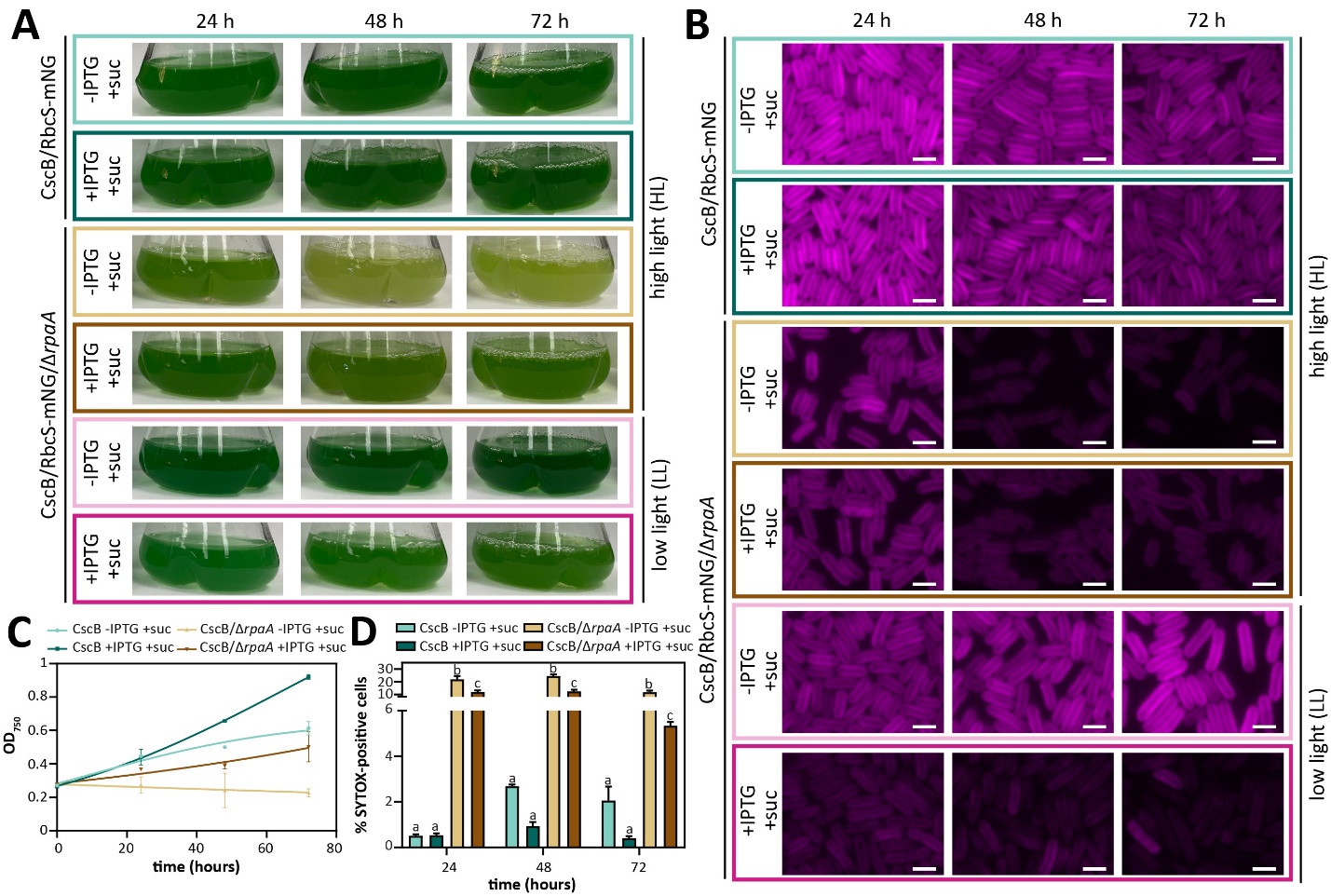

**Supplementary Figure S13. Effect of ambient air conditions on chlorophyll, ROS levels, growth and viability. (A)** Changes in the appearance of the cultures following the transference to ambient air conditions under mixotrophic conditions for the strains CscB/RbcS-mNG and CscB/RbcS-mNG/Δ*rpaA*. **(B)** Chlorophyll changes of the strains in response to ambient air conditions under mixotrophic conditions for the strains CscB/RbcS-mNG and CscB/RbcS-mNG/Δ*rpaA* by tracking the red channel. Scale bar: 2 µm. **(C)** Growth curves of cultures of the strains CscB/RbcS-mNG and CscB/RbcS-mNG/Δ*rpaA* following the transference to ambient air conditions under mixotrophic conditions. Averages of ≥3 independent biological replicates are shown ± SD. **(D)** Percentage of SYTOX-positive cells (dead cells) following the transference to ambient air conditions under mixotrophic conditions for the strains CscB/RbcS-mNG and CscB/RbcS-mNG/Δ*rpaA*. **(C, D)** Averages of ≥3 independent biological replicates are shown + SD. Significance was calculated by one-way ANOVA followed by Tukey’s multiple comparison test. Data points labeled with different letters are significantly different (*P* < 0.05).

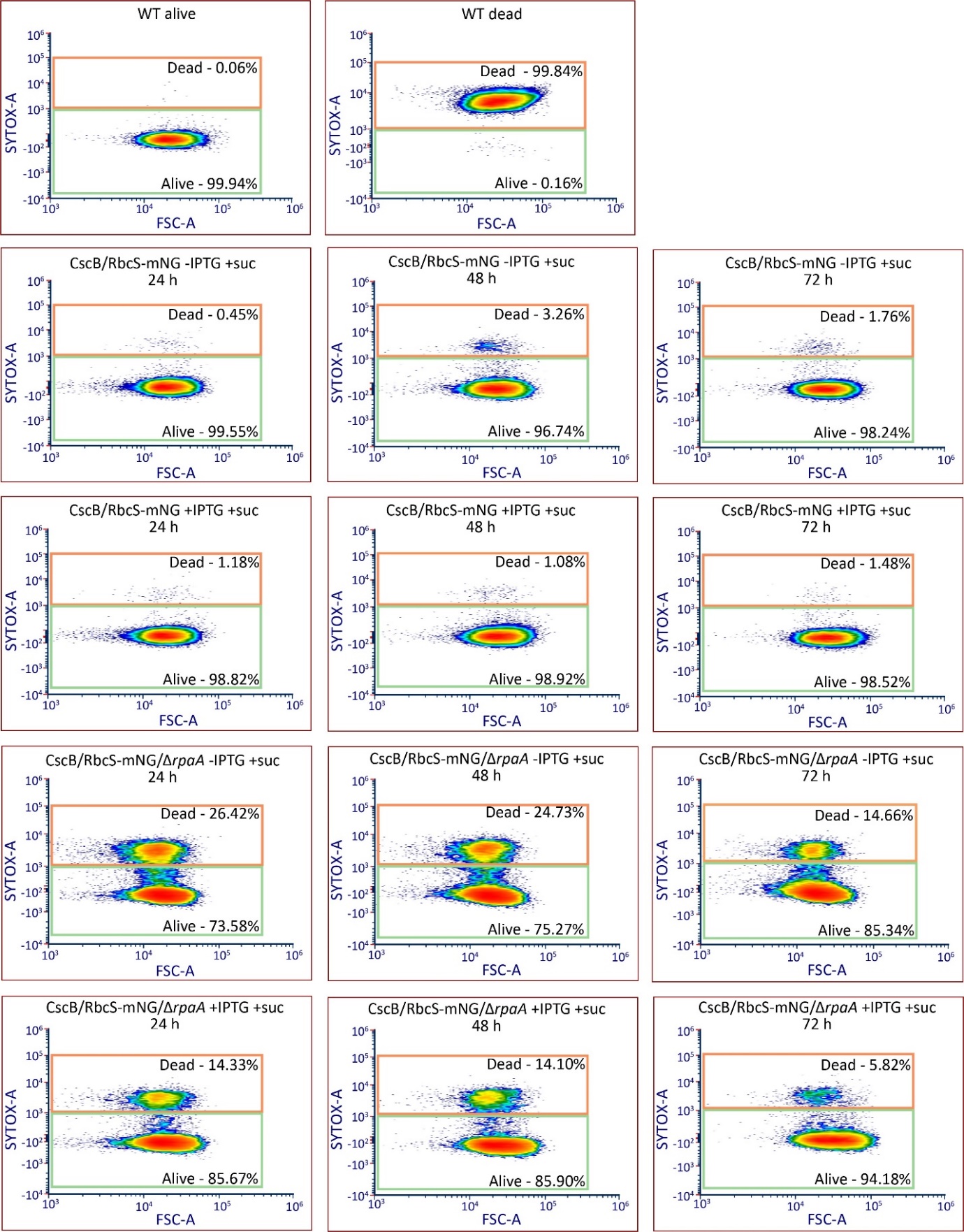

**Supplementary Figure S14. Dot plots of SYTOX blue flow cytometry after CO_2_ down-shift revealed a negative in the viability of Δ*rpaA* mutant even in absence of sucrose feeding.** Alive, SYTOX-negative cells; Dead, SYTOX-positive cells; FSC-A, forward scatter area; SYTOX-A, SYTOX blue area.
